# Human Th17- and IgG3-associated autoimmunity induced by a translocating gut pathobiont

**DOI:** 10.1101/2023.06.29.546430

**Authors:** Konrad Gronke, Mytien Nguyen, Noemi Santamaria, Julia Schumacher, Yi Yang, Nicole Sonnert, Shana Leopold, Anjelica L. Martin, Remy Hallet, Kirsten Richter, David A. Schubert, Guillaume M. Daniel, David Dylus, Marianne Forkel, Silvio Manfredo Vieira, Dorothee Schwinge, Christoph Schramm, Kara G. Lassen, Luca Piali, Noah W. Palm, Christoph Bieniossek, Martin A. Kriegel

**Author notes:** Correspondence (M.A.K.). These authors contributed equally (Konrad Gronke, Mytien Nguyen). Senior authors (Noah W. Palm, Christoph Bieniossek, Martin A. Kriegel).

## Abstract

Extraintestinal autoimmune diseases are multifactorial with translocating gut pathobionts implicated as instigators and perpetuators in mice. However, the microbial contributions to autoimmunity in humans remain largely unclear, including whether specific pathological human adaptive immune responses are triggered by such pathobionts. We show here that the translocating pathobiont *Enterococcus gallinarum* induces human IFNγ^+^ Th17 differentiation and IgG3 subclass switch of anti-*E. gallinarum* RNA and correlating anti-human RNA autoantibody responses in patients with systemic lupus erythematosus and autoimmune hepatitis. Human Th17 induction by *E. gallinarum* is cell-contact dependent and involves TLR8-mediated human monocyte activation. In murine gnotobiotic lupus models, *E. gallinarum* translocation triggers IgG3 anti-RNA autoantibody titers that correlate with renal autoimmune pathophysiology and with disease activity in patients. Overall, we define cellular mechanisms of how a translocating pathobiont induces human T- and B-cell-dependent autoimmune responses, providing a framework for developing host- and microbiota-derived biomarkers and targeted therapies in extraintestinal autoimmune diseases.

**One Sentence Summary:** Translocating pathobiont *Enterococcus gallinarum* promotes human Th17 and IgG3 autoantibody responses linked to disease activity in autoimmune patients.

## Introduction

Human T helper cell subsets and autoantibodies are critical effectors of autoimmunity. Human Th17 cells have been identified as crucial cell types in a variety of autoimmune diseases as well as in host protection from pathogens (reviewed in (*1*)). Pathogen-induced human Th17 cells produce pro- and anti-inflammatory cytokines depending on the pathogen and local environment (*2*). Selected murine gut bacteria can induce Th17-dependent autoimmune responses in animal models (reviewed in (*3*)). However, little is known about the nature and mechanisms of gut pathobionts on human Th17 induction and autoimmune diseases. A model pathobiont in mice known to induce mucosal Th17 cells in the small intestine are segmented filamentous bacteria (SFB) (*4*). SFB colonization has been linked to extraintestinal autoimmune diseases such as arthritis and lupus in mice (*5*, *6, 7*); however, human Th17-inducing pathobionts remain elusive, in part because of host-specific adaptation of pathobionts such as SFB (*8*). Systematic screening of human microbiota in gnotobiotic mouse models has revealed candidates that induce or inhibit intestinal Th17 cells in mice (*9, 10, 11, 12, 13*). Also, microbiota-reactive Th17 cells have been described in human subjects (*14, 15, 16, 17*), but the mechanisms how gut microbiota induce human Th17 cells and contribute to extraintestinal autoimmunity in patients remain unclear.

A plausible mechanism of how commensal bacteria in the gut may trigger extraintestinal autoimmune diseases is translocation across the gut barrier, followed by direct interactions with host immune cells in secondary lymphoid organs. We have previously shown that a gut pathobiont, *Enterococcus gallinarum* (EG), translocates to mesenteric lymph nodes (MLN), liver, and spleen in autoimmune-prone animals (*18, 19*). *E. gallinarum* translocation led to Th17 and autoantibody responses outside the gut and murine autoimmune pathology related to SLE and autoimmune hepatitis (AIH). DNA from this pathobiont was identified in liver biopsies from SLE and AIH patients (*18*) and a human isolate promotes murine MLN Th17 in a mouse model of primary sclerosing cholangitis (PSC) together with other pathobionts (*20*).

The pathophysiology of both SLE and AIH involves Th17-mediated autoimmunity and systemic antinuclear antibody production (*21, 22, 23*). Th17 cells are known to promote germinal center reactions and autoantibody production by B cells (*24, 25, 26, 27, 28, 29*), linking cellular and humoral autoimmune responses. To date, it is not known how gut pathobionts induce human Th17- and autoantibody-dependent autoimmunity and which autoantibody subclasses are involved in the pathophysiology. Here, we show that the translocating gut pathobiont *E. gallinarum* robustly induces human Th17 differentiation via a contact-dependent mechanism involving TLR8 signals in human monocytes, a process that is blocked by the disease-modifying antirheumatic drug hydroxychloroquine (HCQ). We further demonstrate that *E. gallinarum* promotes switching to human IgG3 subclass production *in vitro* as well as murine IgG3 autoantibody production and autoimmune pathology in gnotobiotic models *in vivo*. Finally, we show that human IgG3 autoantibodies against human RNA and anti-*E. gallinarum* RNA responses are linked in SLE and AIH patients, respectively. In summary, we provide a conceptual framework on how a translocating gut pathobiont induces pathogenic human Th17 and autoantibody responses with broad implications for human extraintestinal autoimmune diseases.

## Results

### *E. gallinarum* stimulates robustly human Th17 responses *in vitro*

To recapitulate the physiological differentiation of human T cells *in vitro*, we stimulated peripheral blood mononuclear cells (PBMCs) from healthy human donors with bacterial lysates from commensal bacteria. We have chosen a previously described *E. gallinarum* isolate derived from an autoimmune-prone murine model that is closely related to human isolates and was shown to translocate from the small intestine to lymphoid organs (*18, 19*). We compared the effects of *E. gallinarum* to the stimulation with an *Enterococcus faecalis* (*E. faecalis,* EF) strain, which is a pore-forming toxin-producing opportunistic pathogen in the same genus, that can also reside in the human intestine (*30*). As an additional enteric bacterium, we chose an *Escherichia coli* (*E. coli,* EC) isolate, which colonizes the human colon, and can also translocate and cause toxin-dependent colonic inflammation and diarrhea (*31*). All three isolates were able to disrupt a human intestinal epithelial cell layer in two *in vitro* models (fig. S1A).

Exposing PBMCs to bacterial lysates led to activation of human T cells that produced several pro-inflammatory cytokines (fig. S1B). Stimulation of anti-CD3/CD28 activated T cells with *E. gallinarum* led to a vigorous induction of IL-17-producing CD4^+^ T cells (Fig. 1, A to C). We also observed an increase in IFNγ-producing CD4^+^ T cells after exposure to *E. gallinarum* lysate (fig. S1, C and D), thus demonstrating that *E. gallinarum* vigorously induces human Th17/Th1 responses, which are known to be pathogenic in autoimmune disorders (*32, 33, 34*). *In vitro*, the genus-related *E. faecalis* lysate also induced IL-17 and IFNγ production, suggesting the molecular triggers leading to this immune profile could be conserved among several *Enterococcus* species similar to certain immunologically active toxins (*30*) or enzymes (*35*). In contrast, *E. coli* lysate exposure was not able to increase IL-17 production significantly (Fig. 1, A to C).

**Fig 1.**
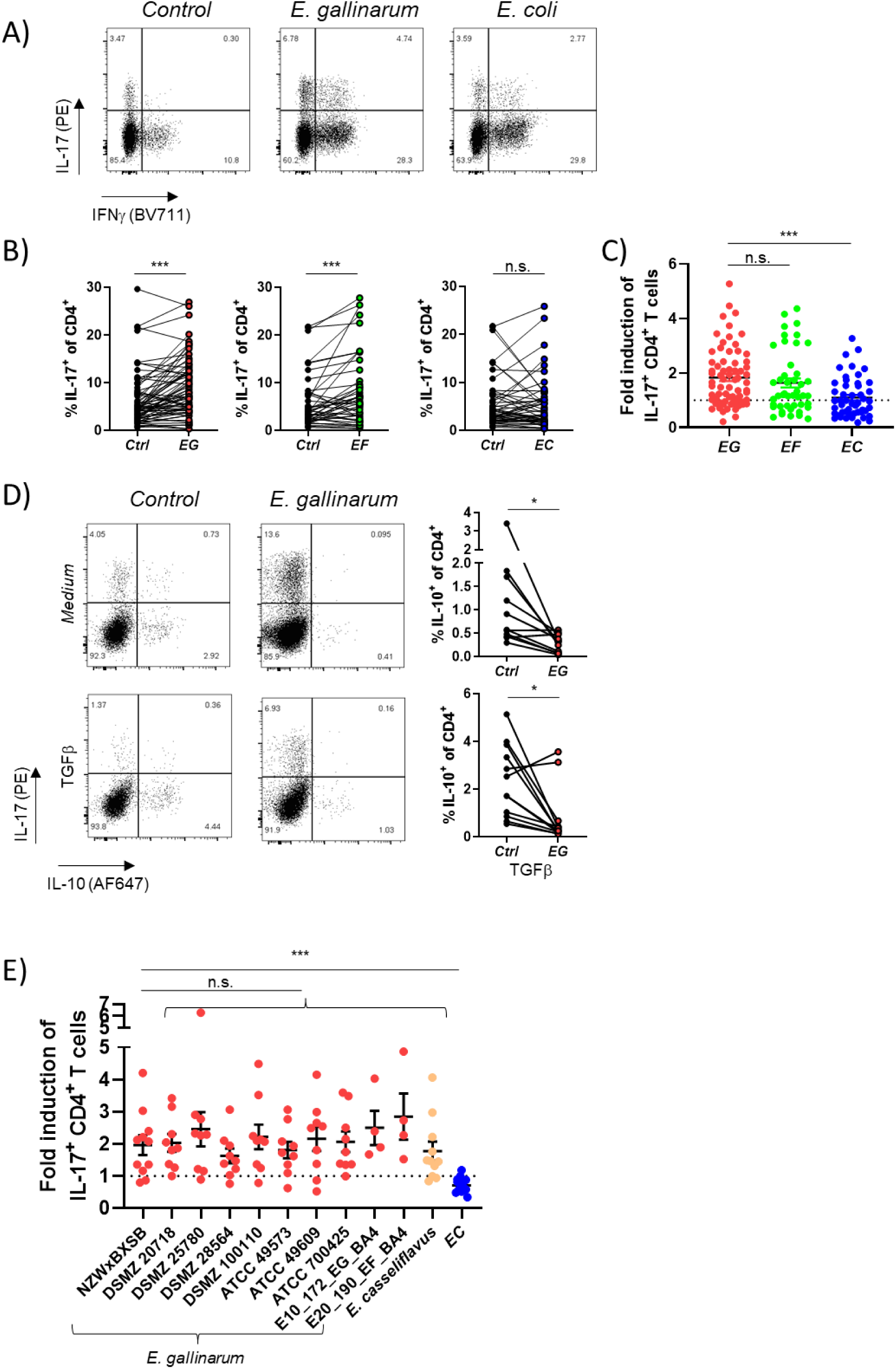
*E. gallinarum* induces IL-17 in human immune cells. Human PBMCs were stimulated with bacterial lysates in the presence of CD3/CD28 activation. Cytokine production by CD4^+^ T cells was assessed by flow cytometry on day 7. a) Representative plots gated on CD4^+^ T cells for no bacterial lysates (*Control*), *E. gallinarum,* and *E. coli* lysate. b) Quantification of IL-17 expression by CD4^+^ T cells in response to bacterial stimulation. c) Fold induction of IL-17 over no bacterial lysates (control) by *E. gallinarum (EG)*, *E. faecalis (EF)* or *E. coli (EC)* in PBMCs on day 7 (EG n=69, EF n=45, EC n=49). d) TGFβ was added to PBMC cultures. Representative plots gated on CD4^+^ T cells for IL-17 and IL-10 production. Reduction of IL-10 production by CD4^+^ T cells in the presence of *E. gallinarum (EG)* lysates without (upper panel) or with (lower panel) additional TGFβ. e) Comparison of several *E. gallinarum* isolates, *E. casseliflavus*, and *E. coli* for fold induction of IL-17^+^ CD4^+^ T cells. Statistical analysis: Unpaired t-test (c,e); paired t-test (b,d)

We also investigated whether bacterial lysates were sufficient to induce human T cell activation without additional T cell receptor (TCR) activation. Again, lysates of *E. gallinarum* and *E. faecalis* alone initiated a strong Th17 effector response, while *E. coli* only induced Th1 cells *in vitro* (fig. S1, E and F).

T cells are highly heterogeneous in composition and types of cytokine responses (*2, 36, 37, 38*), which might determine their contributions to chronic inflammation. Screening cytokine-producing CD4^+^ T cells from PBMCs of healthy subjects *ex vivo*, we observed a wide range of baseline cytokine production for IL-17 and IFNγ (Fig. 1B). Enterococci increased the percentage of IL-17 producing CD4^+^ T cells in almost all donors, regardless of their baseline level of cytokine production (Fig. 1B).

Since mucosally adapted *E. gallinarum* will first encounter immune cells in the lamina propria (*19*), which is an environment that promotes tolerogenic responses, we tested how CD4^+^ T cells would respond to *E. gallinarum* stimulation in the presence of the tolerance-promoting factor TGFβ. While TGFβ alone expectedly increased the percentage of IL-10-producing CD4^+^ T cells, the presence of *E. gallinarum* lysates abolished CD4^+^ T cell-derived IL-10 in almost all donors and led to a robust increase in IL-17^+^ cells (Fig. 1D).

As the tested *E. gallinarum* isolate had been derived from a murine model of autoimmunity, we investigated whether the Th17-dominated immune activation was a conserved feature across other *E. gallinarum* strains, including human-derived isolates we previously showed to share genetic characteristics (*19*). We acquired several *E. gallinarum* isolates derived from humans, mice, and birds, as well as a closely related strain of *Enterococcus casseliflavus*. All tested isolates induced a strong Th17 response in human PBMCs (Fig. 1E). In conclusion, *E. gallinarum* and related strains potently activate human Th17 responses and can overcome mechanisms of immune tolerance.

### *E. gallinarum* leads to the secretion of Th17-polarizing cytokines

Th17 differentiation requires cytokines to be released by surrounding cells in the tissue including myeloid cells. To test whether *E. gallinarum* promotes Th17-skewing conditions, we measured secreted cytokines in the supernatant of PBMC cultures. In the presence of *E. gallinarum* lysates, we observed increased levels of IL-23, IL-6, IL-1β, and IL-1α (Fig. 2A), which are known drivers of Th17 differentiation (*1*). In addition, we observed increased levels of other proinflammatory cytokines such as IFNγ, IL-12, and TNFα (fig. S2A), confirming the broad proinflammatory potential of immune cells activated by *E. gallinarum*. We could also detect an increase in IL-22, which is secreted by cells of the Th17 lineage (*1, 39*). While we had observed a decrease in IL-10 production by T cells after contact with *E. gallinarum* (Fig. 1D), we saw a trend towards higher total cytokine levels in the PBMC supernatant (fig. S2A). The likely source of the majority of IL-10 in this setting is myeloid cells, which are known to release IL-10 in response to bacterial stimuli (*40*).

**Fig 2.**
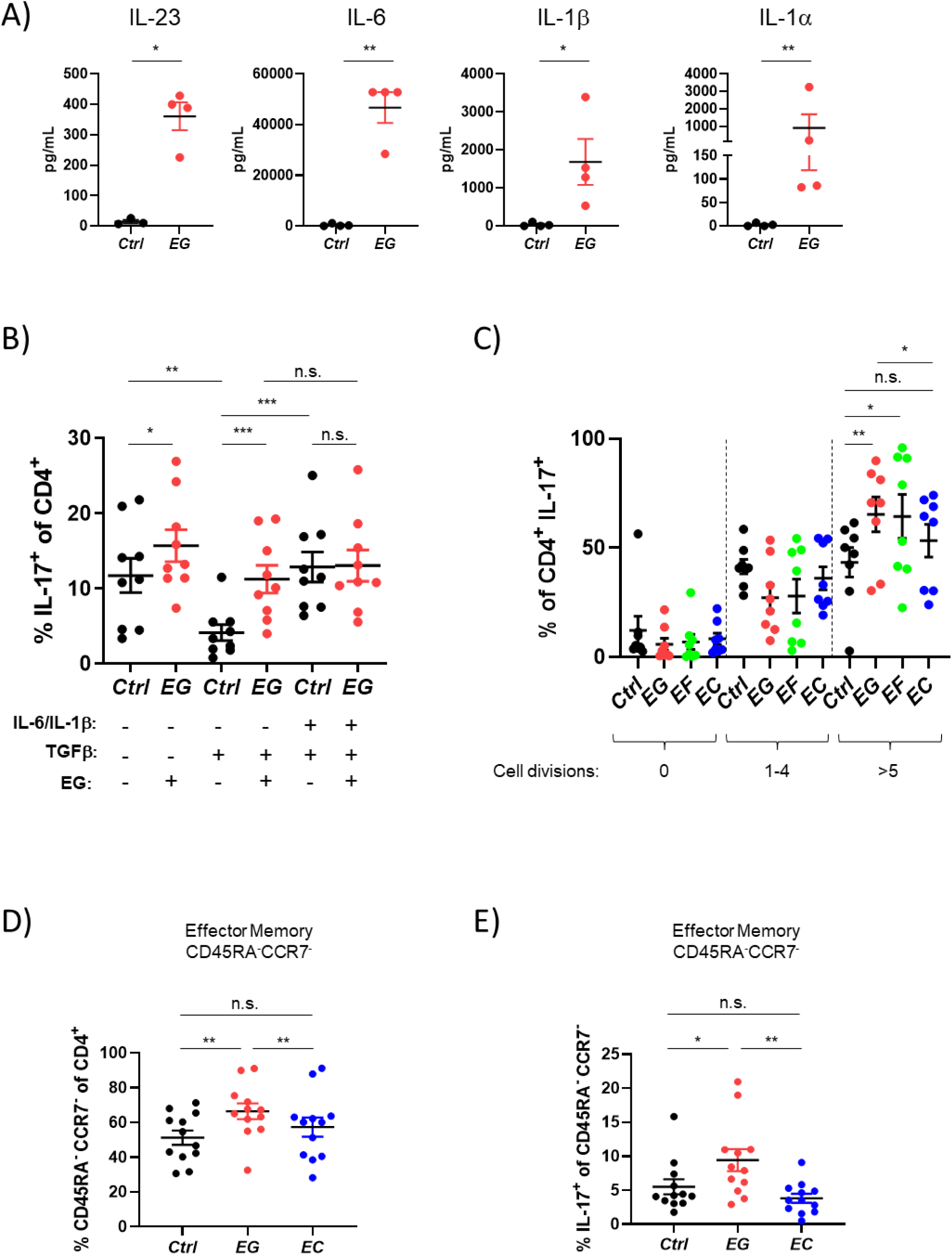
Increase of IL-17 is related to differentiation and proliferation of Th17 cells. Human PBMCs were stimulated with bacterial lysates in the presence of CD3/CD28 activation and analyzed on day 7 after stimulation. a) Secreted proteins were measured in the supernatants by a bead-based multiplex assay after stimulation with *E. gallinarum (EG)* lysate. b) TGFβ and IL-6/IL-1β were added to selected cultures. Percentage of IL-17-producing CD4^+^ T cells after stimulation with *E. gallinarum (EG)*. c) PBMCs were incubated with a cell proliferation dye prior to stimulation with *E. gallinarum (EG)*, *E. faecalis (EF)* or *E. coli (EC)* lysate. The number of cell divisions was assessed after one week and shown here for IL-17^+^ CD4^+^ T cells. These were grouped accordingly into cells that had not divided, had undergone 1 to 4 division, and cells that had divided more than 5 times. d) Percentage of CD45RA^-^CCR7^-^ effector memory cells of CD4^+^ T cells after stimulation with *E. gallinarum (EG)* or *E. coli (EC)*. e) Percentage of IL-17^+^ cells of CD4^+^CD45RA^-^CCR7^-^ effector memory T cells after stimulation with *E. gallinarum (EG)* or *E. coli (EC)*. Statistical analysis: paired t-test (b,c,d,e), Ratio paired t-test (a)

Next, we wanted to test if *E. gallinarum* stimulation provides synergistic effects for Th17 differentiation beyond classical cytokine-induced differentiation. After adding TGFβ to the *in vitro* cultures without *E. gallinarum*, we observed a decrease in baseline IL-17 production. This decrease was entirely overcome by the presence of *E. gallinarum* lysates (Fig. 2B). By providing recombinant IL-6 and IL-1β to cultures already containing TGFβ, we recreated optimal conditions for Th17 differentiation *in vitro*. *E. gallinarum* did not lead to a synergistic increase in IL-17-producing CD4^+^ T cells beyond these Th17-skewing cytokines (Fig. 2B). This finding indicates that *E. gallinarum* induces classical Th17 differentiation programs.

### IL-17^+^ effector memory T cells proliferate in response to *E. gallinarum*

Proliferation is necessary for differentiation of naive T cells into effector lineages (*41*) but may also lead to the specific expansion of pre-differentiated cells. To assess the role of proliferation in our system, we analyzed the number of divisions with a cell tracer. We observed an increase in the proliferation of CD4^+^ T cells after they have been stimulated with any of the bacterial lysates (fig. S2, B and C). Focusing on IL-17-producing CD4^+^ T cells, we observed that CD4^+^ T cells that had been in contact with *E. gallinarum* or *E. faecalis* had undergone significantly more rounds of proliferation compared to unexposed or *E. coli*-stimulated T cells (Fig. 2C). The proliferative effects on Th17 cells were again selective for enterococci as all bacteria, including *E. coli,* induced vigorous proliferation of IFNγ-secreting cells (fig. S2, B and C).

Next, we analyzed the T cell effector subtype distribution given that effector memory T cells can induce innate inflammation and autoimmune pathology (*42*). After *E. gallinarum* stimulation, we observed significantly more CD45RA^-^CCR7^-^ “effector memory” T cells compared to controls and *E. coli* stimulation, respectively (Fig. 2D and fig. S2D). Accordingly, there was a decrease in CD45RA^+^CCR7^-^ “terminally differentiated” cells (Temra) (fig. S2E). These effector memory-like T cells contained a higher percentage of IL-17-producing CD4^+^ T cells in response to *E. gallinarum* (Fig. 2E and fig. S2F). We conclude that *E. gallinarum* preferentially leads to the expansion of Th17 cells with effector functions.

### Cell-cell contact between antigen presenting cells (APCs) and T cells is necessary for Th17 induction

Bacterial products can activate T cells in a TCR-independent manner by directly acting on T cells or by inducing “bystander activation” via cytokines derived from the surrounding environment. Bystander activation has been observed during infections (*43*) as well as host-microbiota interactions (*44*). Bystander activation has also been well described in autoimmune conditions and does not require specific antigen presentation by APCs in contrast to classical antigen-dependent activation (*45, 46*), which can trigger cross-reactive T cells (*3*).

To elucidate whether *E. gallinarum* could directly activate T cells, we purified CD4^+^ T cells from PBMCs and exposed them to lysates without APCs. We found that *E. gallinarum* stimulation did not lead to a meaningful increase in Th17 cells in the absence of APCs (Fig. 3A).

**Fig 3.**
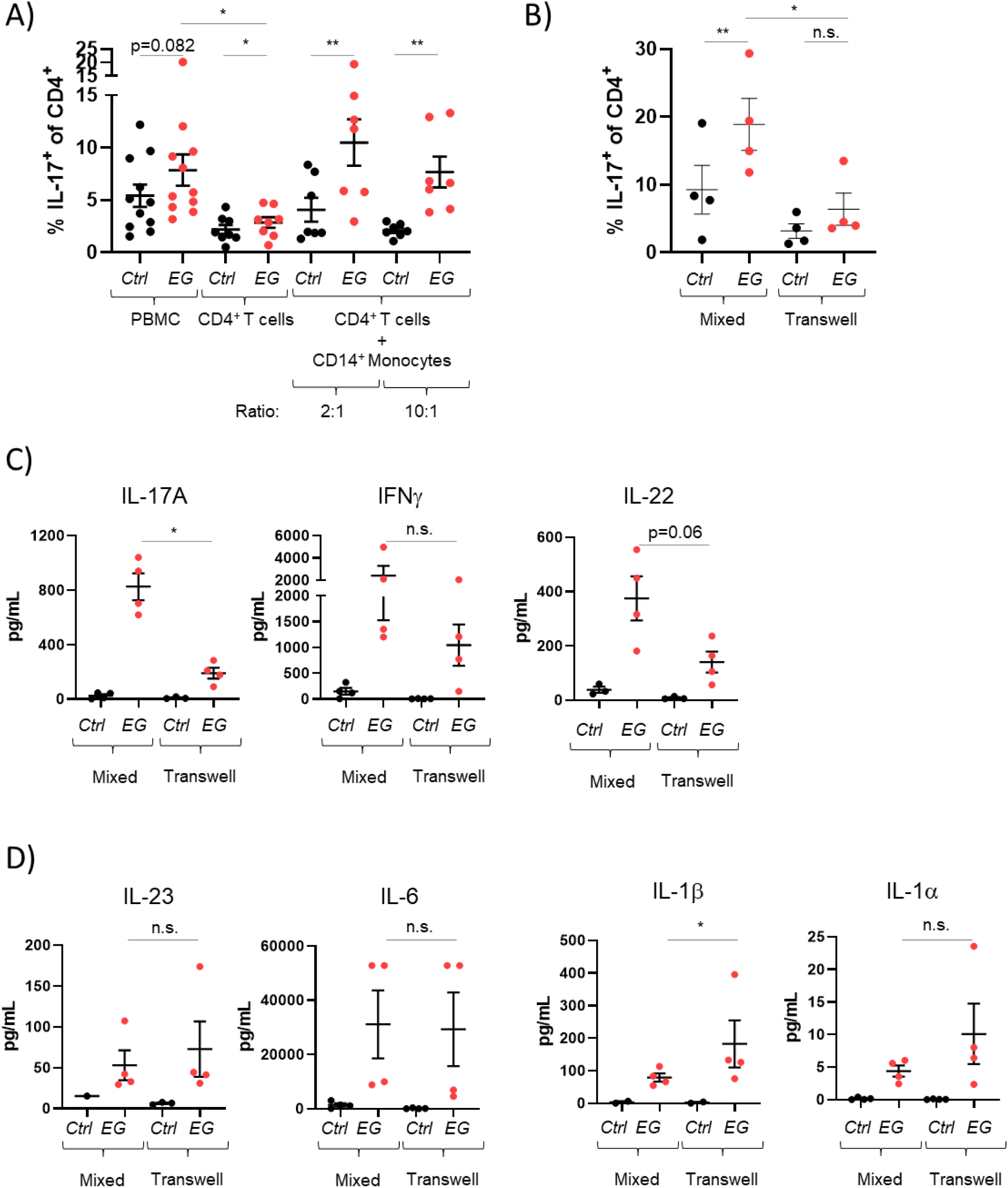
IL-17 induction by *E. gallinarum* depends on direct cell-cell contact between human CD4^+^ T cells and monocytes. Human PBMCs were prepared and CD14^+^ monocytes and CD4^+^ T cells were isolated via magnetic bead sorting. Cells were stimulated with *E. gallinarum (EG)* lysate for 7 days in the presence of CD3/CD28 activation. a) PBMCs, isolated CD4^+^ T cells or combined cultures of CD4^+^ T cells and CD14^+^ monocytes (in two different ratios of CD4^+^ T cells exceeding CD14^+^ monocytes) were stimulated with *E. gallinarum (EG)* and cytokine production by CD4^+^ T cells was assessed by flow cytometry. b) CD4^+^ T cells and CD14^+^ monocytes were either mixed or kept separated by a transwell system and stimulated with *E. gallinarum (EG)* lysates. The percentage of IL-17-producing CD4^+^ T cells was assessed by flow cytometry. c,d) Secreted proteins from b) were measured in the supernatant by a bead-based multiplex assay. Statistical analysis: paired t-test (a,b), Ratio paired t-test (c,d)

Next, we sought to determine whether cell-cell contact between APCs and T cells was necessary or whether cytokine production alone, as seen in bystander activation, would be sufficient to induce Th17. We isolated human CD14^+^ monocytes, which are capable of antigen presentation as well as secretion of T cell-differentiating cytokines. Monocytes from the same donor were co-cultured with the corresponding CD4^+^ T cells in different ratios. When CD14^+^ monocytes and T cells were able to interact directly, *E. gallinarum* lysate induced strong IL-17 responses (Fig. 3A).

To differentiate between the effects of cytokines and surface receptors expressed by APCs, we separated purified CD4^+^ T cells and CD14^+^ monocytes in a transwell system that does not allow cell-cell contact while enabling exchange of soluble mediators. When CD4^+^ T cells and CD14^+^ monocytes were physically separated, we observed a significantly reduced percentage of IL-17-producing CD4^+^ T cells after bacterial stimulation (Fig. 3B) despite additional TCR activation by anti-CD3/-CD28 beads. We further analyzed secreted cytokines in the culture supernatants and detected a decrease in IL-17 and IL-22 in all donors, confirming diminished Th17 differentiation without direct APC-T cell contact (Fig. 3C).

Notably, we did not observe a significant decrease in IFNγ production nor Th1-promoting IL-12 by separating APCs and T cells, thus demonstrating that contact dependency was preferentially required for Th17, but less so for Th1 differentiation induced by bacteria (Fig. 3C and fig. S3A).

Finally, we analyzed Th17-promoting soluble mediators induced by *E. gallinarum* (IL-23, IL-6, IL-1β, IL-1α). We did not observe a reduction of these cytokines after transwell separation, emphasizing the need for an additional contact-dependent co-stimulatory factor for Th17 polarization (Fig. 3D).

Taken together, Th17 differentiation by *E. gallinarum* requires not only the secretion of T cell-skewing cytokines, but also the interaction of surface molecules between T cells and APCs.

### *E. gallinarum* induces the expression of co-stimulatory molecule CD86 on the surface of APCs

The nature of the interaction between APCs and lymphocytes is defined by the expression and type of costimulatory molecules (*47*). We hypothesized that *E. gallinarum* induces molecules on the surface of APCs that facilitate a productive interaction with T cells leading to their activation. After stimulation of PBMCs with the different bacterial lysates, CD86 expression was increased on the surface of CD14^+^ monocytes in response to enterococci but not *E. coli* (Fig. 4A). Testing surface expression of the closely related costimulatory molecule CD80, we detected a slight increase induced by enterococci and a much stronger increase induced by *E. coli* lysates (Fig. 4A).

**Fig 4.**
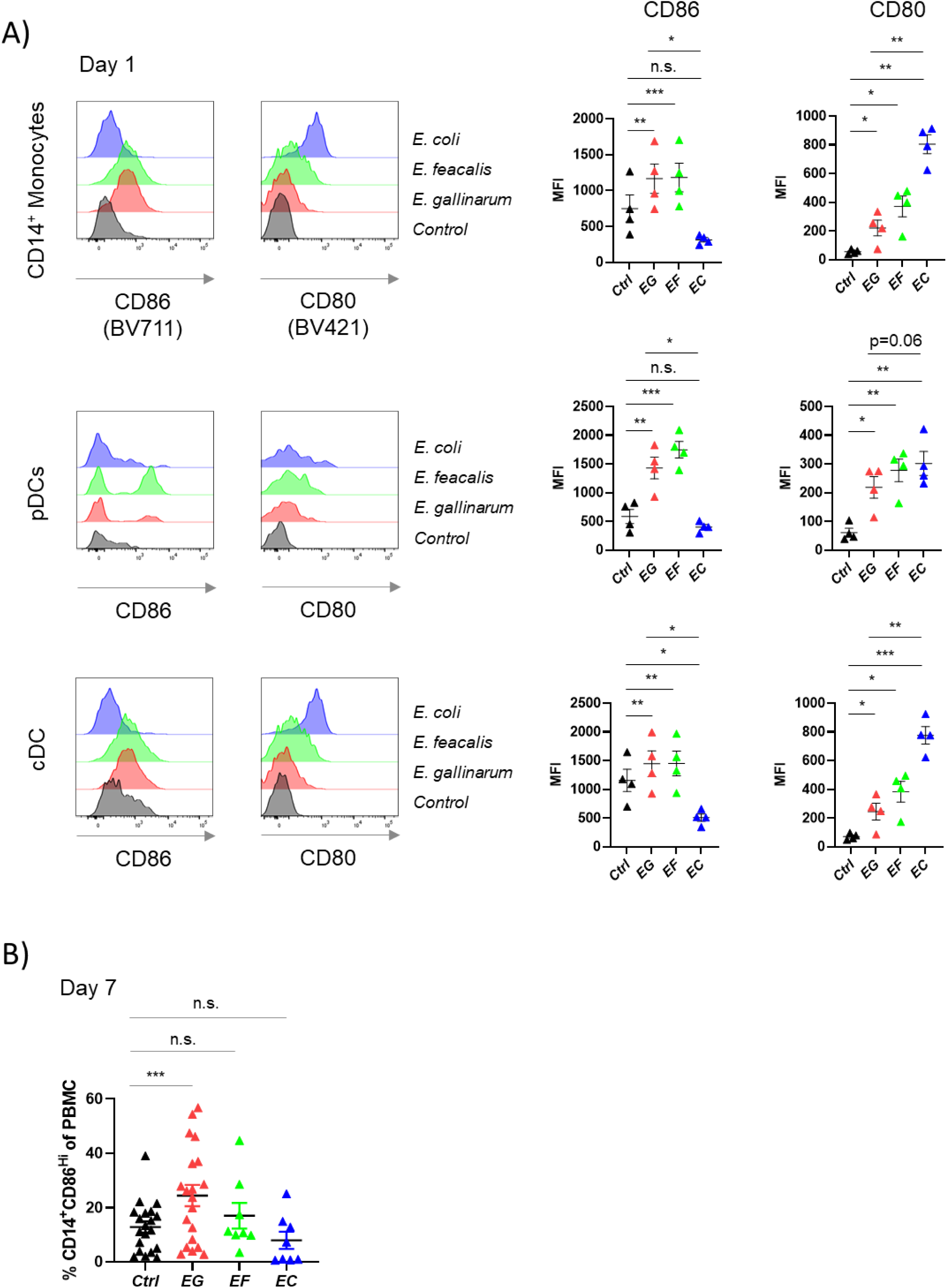
*E. gallinarum* upregulates specific co-stimulatory molecules on APCs and leads to TCR activation in PBMCs. Human PBMCs were stimulated with *E. gallinarum (EG)*, *E. faecalis (EF)*, or *E. coli (EC)* lysates. a) Expression of co-stimulatory molecules CD80 and CD86 by different APCs assessed by flow cytometry on day 1 after stimulation. b) Percentage of CD14^+^CD86^High^ leucocytes on day 7 after bacterial stimulation. Statistical analysis: paired t-test (a,b)

We also observed a similar pattern of strong CD86 induction by enterococci and CD80 induction by *E. coli* in plasmacytoid dendritic cells (pDC) and classical dendritic cells (cDC), which have both been implicated in autoimmune pathologies (*48*) (Fig. 4A). On the other hand, there was no induction of CD86 or CD80 on human B cells and CD16^+^ monocytes, respectively (fig. S4A).

The expression of CD86 on CD14^+^ monocytes was robust and long-lasting *in vitro* and was sustained in cultures after seven days, indicating ongoing positive feedback mechanisms (Fig. 4B). A role for bidirectional effects of activated T cells in maintaining CD86 expression on APCs was supported by the finding that activation of T cells by anti-CD3/CD28 was also able to induce CD86 expression on CD14^+^ monocytes in culture, whereas the presence of *E. coli* prevented this effect (fig. S4B). Overall, these findings demonstrate that bacterial lysates induce specific patterns of costimulatory receptors on the surface of APCs, which facilitate sustained interactions with T cells.

To determine if this interaction would favor the expansion of single T cell clones, we stained for the distribution of several TCR α and β chains before and after cultivation in the presence of bacterial lysates. If pre-existing *Enterococcus*-specific T cell clones would expand, they would be expected to dominate the T cell repertoire in the PBMC cultures. We did not observe any major differences in the TCR repertoire of CCR6^+^ CD4^+^ T cells, which are enriched for Th17 cells, compared to the total CD4^+^ T cell population in PBMC of healthy subjects (fig. S4C). However, the distribution of TCR chains changed during seven days of culture in the presence or absence of bacterial lysates with specific populations displaying growth advantages. Focusing specifically on IL-17- and IFNγ-producing CD4^+^ T cells, we detected an overrepresentation of T cells with certain TCR chains, suggesting that not all CD4^+^ T cells contribute equally to cytokine production after bacterial challenge. However, there were no differences in TCR chain distributions between cultures that had been stimulated with either *E. gallinarum, E. faecalis* or *E. coli* (fig. S4C). Therefore, we conclude that the overrepresentation of certain TCR chains among cytokine-producing T cells was not the result of expansion of antigen-specific clones, but rather represent a T cell fraction that was poised to produce large amounts of cytokines after activation by gut bacteria.

In summary, a contact-dependent APC-T cell interaction induced by *E. gallinarum* leads to Th17 differentiation, but not to an outgrowth of individual T cell clones.

### An unsecreted component of *E. gallinarum* leads to IL-17 induction via TLR8 activation

After observing a robust Th17 polarization in human PBMC cultures, we wondered how *E. gallinarum* drives this immune activation. To further delineate the mechanism, we first tested if a secreted component of *E. gallinarum* could induce Th17 differentiation. Induction of IL-17-producing CD4^+^ T cells in PBMCs occurred in the presence of *E. gallinarum* lysates and was absent after addition of *E. gallinarum* supernatants (Fig. 5A). Therefore, we next processed the bacterial lysates to deplete potential stimulatory components. Digestion of proteins by proteinase K did not affect Th17 differentiation (Fig. 5B), nor did heat-inactivation of the bacterial lysates reduce the stimulatory capacity of *E. gallinarum* (fig. S5A).

**Fig 5.**
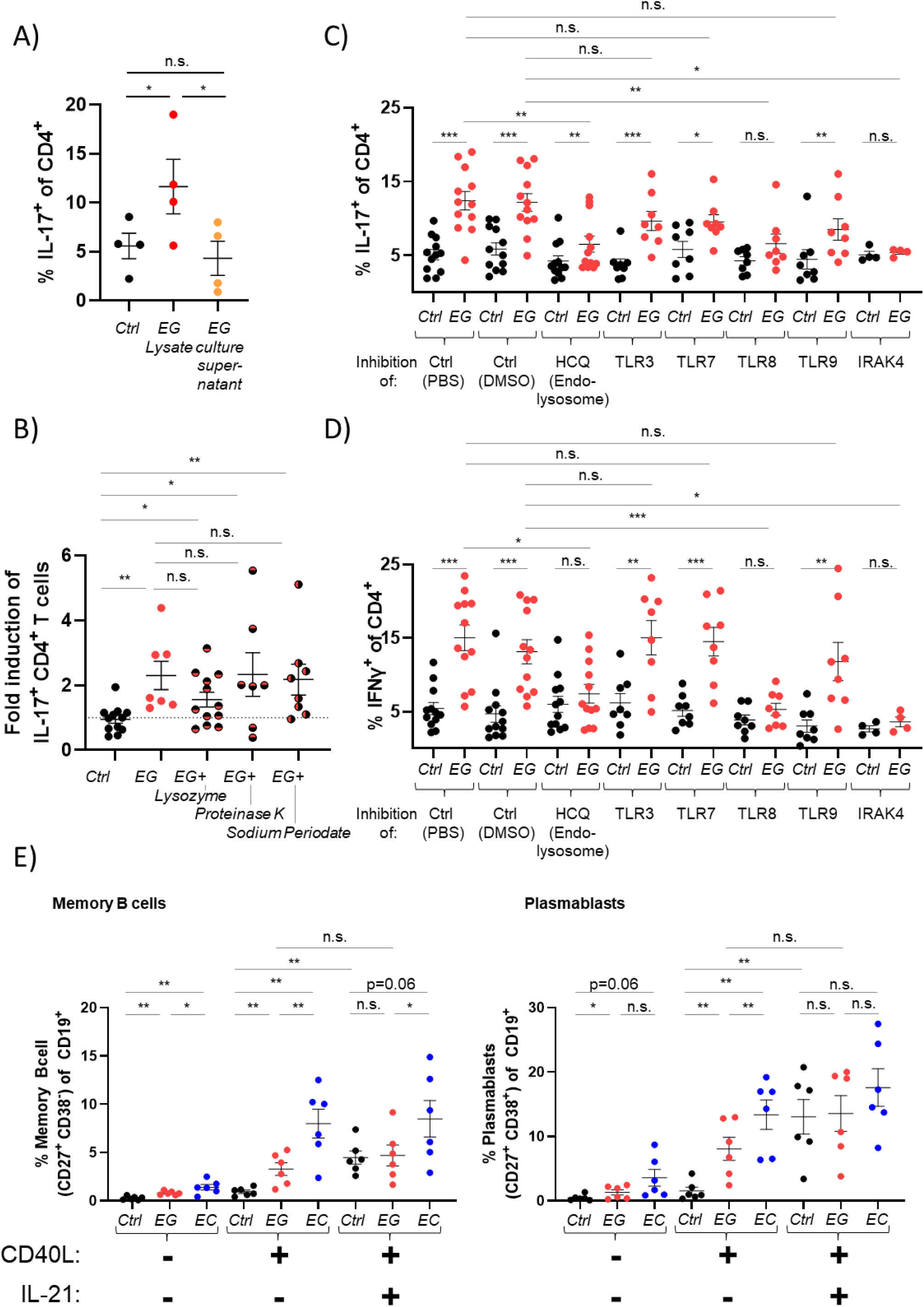
A non-secreted component of *E. gallinarum* activates TLR8 in PBMC leading to IL-17 induction in CD4^+^ T cells. Human PBMCs were stimulated with bacterial lysates in the presence of CD3/CD28 activation. Cytokine production by CD4^+^ T cells was assessed by flow cytometry on day 7. a) Cells were unstimulated (Ctrl) or treated with *E. gallinarum (EG)* lysate or the supernatant of *E. gallinarum* bacterial cultures. b) *E. gallinarum (EG)* lysates were treated with lysozyme, proteinase K, or sodium periodate prior to being added to PBMC cultures. Fold induction of percentage of IL-17-producing CD4^+^ T cells is shown. c,d) PBMCs were treated with soluble inhibitors (hydroxychloroquine, selective TLR3/7/8/9 and IRAK4 inhibitors) of several pattern recognition receptors as indicated. Cells were subsequently stimulated with *E. gallinarum (EG)* lysate. c) IL-17 and d) IFNγ induction in CD4^+^ T cells was measured on day 7 by flow cytometry. e) In addition to stimulation with *E. gallinarum (EG)* or *E. coli (EC)* lysates, PBMCs were treated with recombinant CD40L and IL-21. B cell populations were assessed by flow cytometry on day 7. Statistical analysis: paired t-test (a,c,d,e), unpaired t-test (b)

As we had not observed signs of antigen dependence for Th17 induction, we hypothesized that *E. gallinarum* activates immune cells via pattern recognition receptors (PRRs). Therefore, we inhibited selective PRRs in PBMC cultures prior to stimulation with *E. gallinarum*. We found that neither the inhibition of NOD1 or NOD2 nor TLR2 or TLR5 significantly reduced IL-17 production (fig. S5B). Accordingly, lysozyme, which depletes peptidoglycan, a well-known pro-inflammatory NOD2- and TLR2-ligand, did not affect the capacity of *E. gallinarum* to induce IL-17 responses, nor did the denaturation of carbohydrates by sodium periodate have any effect (Fig. 5B). On the other hand, selective pharmacologic inhibition of IRAK4, a signaling molecule downstream of both TLR and IL-1 signaling, inhibited Th17 differentiation (Fig. 5C), suggesting that additional TLRs are involved in this process.

TLRs can be found on the surface or within the endosomal compartment of cells. Hydroxychloroquine (HCQ) is a disease-modifying antirheumatic drug commonly used to treat SLE patients that acts on endosomal TLRs. It prevents the acidification of the endolysosome and thereby prevents signaling via endosomally localized TLRs (TLR3, 7, 8, 9) as well as proper MHCII loading in professional APCs (*49*). Treatment of PBMCs with HCQ led to a significantly reduced production of IL-17 by CD4^+^ T cells in response to *E. gallinarum* stimulation (Fig. 5C). Next, we used selective TLR inhibitors to delineate whether *E. gallinarum* activates any of the intracellular TLRs. Despite inhibition of TLR3, 7 and 9, CD4^+^ T cells could still respond via IL-17 production. The selective inhibition of TLR8, a receptor that recognizes RNA molecules that represent so-called vita-PAMPs (*50, 51*), blocked IL-17 induction in CD4^+^ T cells (Fig. 5C). HCQ and TLR8 inhibition also reduced IFNγ production, while blockade of TLR3, 7 and 9 did not have a significant effect (Fig. 5D).

Finally, we tested if other Th17-relevant cytokines were also affected by TLR8 blockade. *E. gallinarum* induced IL-21 production in CD4^+^ T cells but was abrogated by inhibition of TLR7 or TLR8, respectively (fig. S5C). Taken together, these data demonstrate that *E. gallinarum* exerts its most significant pro-inflammatory effects by engagement with TLR8.

### *E. gallinarum* supports B cell maturation

As IL-21 is a cytokine involved in B cell maturation and induced by TLR8 via bacterial RNA (*51, 52*), we investigated whether *E. gallinarum* influences B cell populations *in vitro*. We isolated PBMCs from healthy subjects and stimulated them with bacterial lysates. Analyzing B cell populations, there was a small but significant increase in memory B cells and plasmablasts after co-culture with *E. gallinarum* lysate (Fig. 5E and fig. S5D). To support B cell survival in the *in vitro* cultures, we added recombinant CD40L. *E. gallinarum* lysates, together with CD40L, increased memory B cells, plasmablasts and plasma cells, respectively. The increase was even higher in cultures exposed to *E. coli* (Fig. 5E).

As we had seen an induction of IL-21 by *E. gallinarum* (fig. S5C), we explored how this might contribute to B cell maturation, also given that IL-21 signaling cooperates with B cell receptor and CD40 signals in germinal center B cell differentiation (*53*). IL-21 did not increase any further mature B cell populations in conditions that had been stimulated with *E. gallinarum* lysates. However, the combination of CD40L and IL-21 increased B cell populations in the absence of bacterial stimuli to the same extent as was observed with *E. gallinarum* (Fig. 5E and fig. S5D). This indicates that *E. gallinarum* supports B cell maturation either by IL-21 or a factor engaging another B cell-maturing pathway.

These data show that *E. gallinarum* can support human Th17 differentiation and B cell maturation, leading to the generation of potential antibody-producing plasma cells. To test if these cells contribute to autoimmune pathology via the production of autoantibodies, we introduced *E. gallinarum* into gnotobiotic murine models.

### *E. gallinarum* progressively translocates to systemic sites and induces mature plasma cells, autoantibodies, and organ dysfunction

For translating human *in vitro* data to relevant *in vivo* autoimmune pathology, we studied an inducible model of lupus-like disease in gnotobiotic mice. We monocolonized germ-free mice with single bacterial strains and subsequently treated them with topical imiquimod (IMQ) to induce autoimmune pathology (*54, 55*) (fig. S6A).

Analogous to other *E. gallinarum*-driven disease models (*18*), we observed an increase of IL-17-producing CD4^+^ T cells in the mesenteric lymph nodes (MLN) in response to *E. gallinarum* monocolonization (Fig. 6A). Similar to what we had seen with human T cells *in vitro*, *E. coli* did not lead to an increase of IL-17-producing CD4^+^ T cells in MLNs, but there was a trend towards increased Th17 after *E. faecalis* monocolonization (Fig. 6A). Only *E. gallinarum*, but not any of the other gut bacteria, led to splenomegaly in IMQ-treated mice, whereas hepatomegaly was seen in all monocolonized mice (Fig. 6B and fig. S6, B and C). As the spleen is a systemic site, we wondered if bacteria could disseminate to and beyond the “firewall” organs (MLN and liver) once they crossed the intestinal barrier. We could detect viable bacteria of all strains in the MLNs and liver, but only *E. gallinarum* could be cultured in significant amounts from the spleens of treated mice (Fig. 6C). These findings suggest a progressive, time-dependent translocation of *E. gallinarum* from the intestine, which could be confirmed in naturally colonized NZW x BXSB F_1_ SPF mice, a model for lupus-like disease, from which *E. gallinarum* has been originally isolated (*18*) (fig. S6D). In these hybrid mice, naturally colonized *E. gallinarum* crosses the intestinal barrier and colonizes first mesenteric veins and lymph nodes, before migrating further through the liver in order to eventually reach the spleen as a systemic site that is not connected to the enterohepatic circulation (fig. S6D).

**Fig 6.**
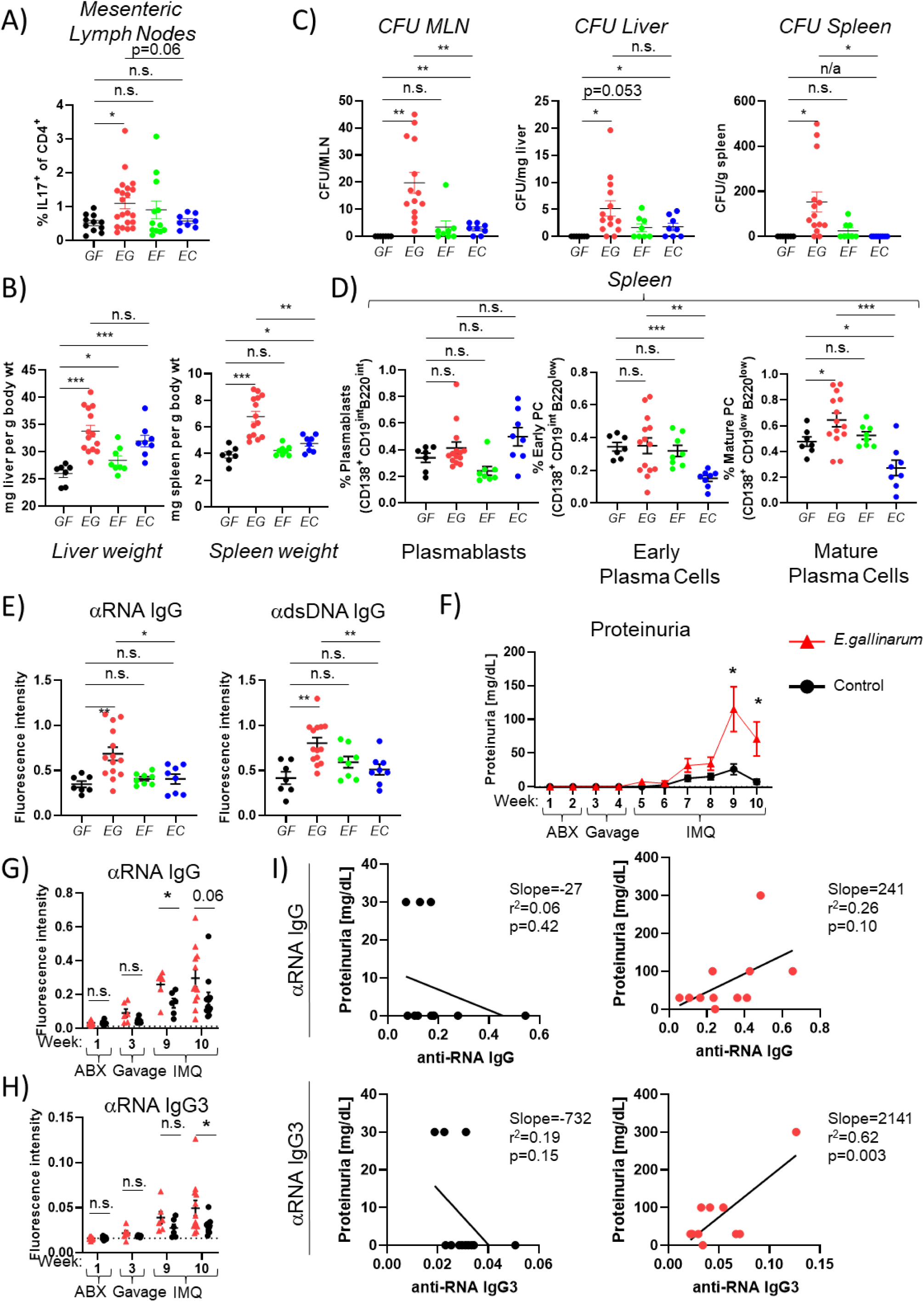
*E. gallinarum* induces extraintestinal Th17- and IgG3-dominated immune responses with mature plasma cell enrichment and lupus-related proteinuria in mice. a-e) C57BL/6-germ-free mice were monocolonized with *E. galllinarum (EG)*, *E. faecalis (EF)* or *E. coli (EC)* and treated with topical imiquimod (IMQ) for 8 weeks. See fig. S6 for schematic outlines of the *in vivo* monocolonization experiments. a) Mesenteric lymph nodes (MLN) were isolated and restimulated in the presence of PMA/ionomycin. IL-17 induction in CD4^+^ T cells was measured by flow cytometry. b) Relative liver (left) and spleen (right) weight of monocolonized mice after 8 weeks of treatment. c) Organs (MLN, liver, spleen) were homogenized and plated on bacterial culture plates. Colony forming units (CFU) were counted and normalized to organ weight. d) Differentiated B cell populations in the spleen of monocolonized mice as determined by flow cytometry. e) Serum reactivity against RNA (left) and dsDNA (right) of monocolonized mice as determined by ELISA. f-i) C57BL/6 SPF mice were treated with oral broad-spectrum antibiotics (ABX) and subsequently gavaged with 2×10^8^ *E. gallinarum* CFUs or PBS (Control). After two weeks of gavage, mice were treated with topical imiquimod every other day as indicated in fig. S6i. Urine and blood were collected. Treatment groups were divided into two cohorts, blood was taken every two weeks, alternating between the cohorts and shifting by a week. See fig. S6i for a schematic outline of the *in vivo* experiments. f) Proteinuria in *E. gallinarum-* vs PBS-gavaged mice measured over the course of the interventions until a total of 10 weeks. g,h) Serum reactivity for IgG (g) and IgG3 (h) against RNA in control and *E. gallinarum* treated mice before ABX treatment (Week 1), after first gavage (Week 3), or after 6 weeks of IMQ treatment (Week 9 and 10). The dotted line represents the detection threshold. i) Correlation between IgG serum reactivity (upper panels) or IgG3 reactivity (lower panels) and proteinuria for control (PBS)- (left) and *E. gallinarum*-gavaged (right) mice. Statistical analysis: Unpaired t-test (a,b,c,d,e,f,g,h); linear regression (i)

Besides an expansion in organ size, we wondered whether the presence of *E. gallinarum* in the spleen would also lead to an altered distribution of B cell populations as we had observed *in vitro*. Indeed, we discovered a significant increase in mature plasma cells in the spleens of mice that had been monocolonized with *E. gallinarum*, but not with *E. faecalis*, whereas *E. coli* colonization unexpectedly reduced plasma cell numbers significantly (Fig. 6D). As mature plasma cells are essential to produce antibodies, including autoantibodies, we next explored if the increase in mature plasma cells would also be reflected in the serum titers of anti-nucleic acid autoantibodies. We detected markedly increased anti-RNA and anti-dsDNA autoantibodies in the serum of *E. gallinarum*-monocolonized mice (Fig. 6E). Increased serum autoantibody titers therefore paralleled increased numbers of mature plasma cells in the spleen. Furthermore, the combination of *E. gallinarum* and IMQ was more potent in inducing systemic autoantibodies than IMQ alone or IMQ plus a resident microbiota from non-autoimmune-prone mice (fig. S6E). Despite the presence of autoantibodies, we did not detect any end-organ damage as indicated by proteinuria at this time point. Thus, we decided to continue the IMQ treatment for 28 weeks for the enterococci-gavaged mice (fig. S6F). After half a year of colonization and IMQ treatment, we detected proteinuria and glomerulonephritis histologically in individual mice but not across the entire animal cohort (fig. S6, G and H), similar to other autoimmune pathobiont monocolonization studies with IMQ (*55, 56*). This variability likely reflects differing within-host evolution in individual mice as shown for *E. gallinarum* and *Lactobacillus reuteri*, another translocating pathobiont (*19*). In addition, lupus-like disease in gnotobiotic models likely requires additional microbiota or other environmental stimuli for full penetrance of disease pathology.

Taken together, our data shows that *E. gallinarum* is a potent inducer of extraintestinal Th17 responses in vivo and subsequently colonizes systemic sites beyond the firewall organs, leading to the expansion of splenic plasma cells and production of autoantibodies.

### *E. gallinarum* colonization aggravates lupus-like renal disease in IMQ-treated SPF mice

To investigate the effects of *E. gallinarum* with additional microbiota signals, we gavaged *E. gallinarum* into an existing microbiota of SPF mice after antibiotic preconditioning followed by topical IMQ treatment (fig. S6, I and J). Mice that had received *E. gallinarum* developed more severe proteinuria reflecting lupus nephritis in this model (Fig. 6F). In addition, the increase in renal dysfunction was paralleled by the induction of anti-RNA and -dsDNA autoantibodies systemically (Fig. 6G and fig. S6K).

Next, we wanted to investigate the IgG subtypes induced by *E. gallinarum* since they may differ in their effector capabilities and disease involvement. T cell-derived cytokines IL-17 and IL-21 are important for immunoglobulin class switching to certain IgG subclasses (*57, 58*). Murine IgG3 is a pathogenic factor for genetic mouse models of autoimmunity (*59, 60, 61*). *E. gallinarum* induced significantly more IgG3 antibodies targeting RNA in IMQ-treated SPF mice (Fig. 6H and fig. S6L). To assess its disease relevance, we correlated anti-RNA IgG and IgG3 titers with the level of proteinuria indicative of glomerular damage. While anti-RNA IgG levels correlated weakly with proteinuria, we observed a robust association between IgG3 titers and the degree of proteinuria (Fig. 6I), suggesting a role for IgG3 in the pathogenesis of lupus-like renal disease. In summary, introducing *E. gallinarum* into a complex microbiota revealed its effects on extraintestinal autoimmune pathology and induction of systemic IgG3 responses that correlated with disease severity.

### IgG3 subclass antibodies from SLE patients react proportionally against *E. gallinarum* and human RNA

Nuclear autoantibodies are commonly found in SLE and AIH patients and contribute to disease pathology. Since *E. gallinarum* influences B cell responses and supports the generation of autoantibodies *in vivo*, we wondered if this would also be reflected in human SLE and AIH patient sera *ex vivo*.

We compared sera from healthy subjects, SLE patients, and AIH patients, respectively, and tested their reactivity against human and bacterial RNA preparations. We chose RNA as a nuclear antigen, as it can act as vita-PAMP and is known to stimulate TLR8 (*50, 51*), a pathway we identified to trigger *E. gallinarum*-induced human Th17 cells *in vitro* (Fig. 5, C and D).

Serum from SLE patients, and to a lesser degree also from AIH patients, reacted strongly against human RNA, confirming the presence of systemic autoantibodies (Fig. 7A). Next, we assessed if SLE and AIH patients responded against *E. gallinarum* RNA. Both patient groups displayed significantly higher IgG titers against *E. gallinarum* RNA than healthy controls (Fig. 7A). The reactivity exhibited a high degree of inter-individual heterogeneity, similar to an SLE cohort in which whole *E. gallinarum* reactivity was demonstrated in a subset of patients with autoreactivity against ribosomal P, an RNA binding protein (*62*). Interestingly, a subgroup of healthy subjects did not contain any antibodies against *E. gallinarum* (Fig. 7A), while all patients exhibited at least some reactivity against the exposed RNA-associated epitopes.

**Fig 7.**
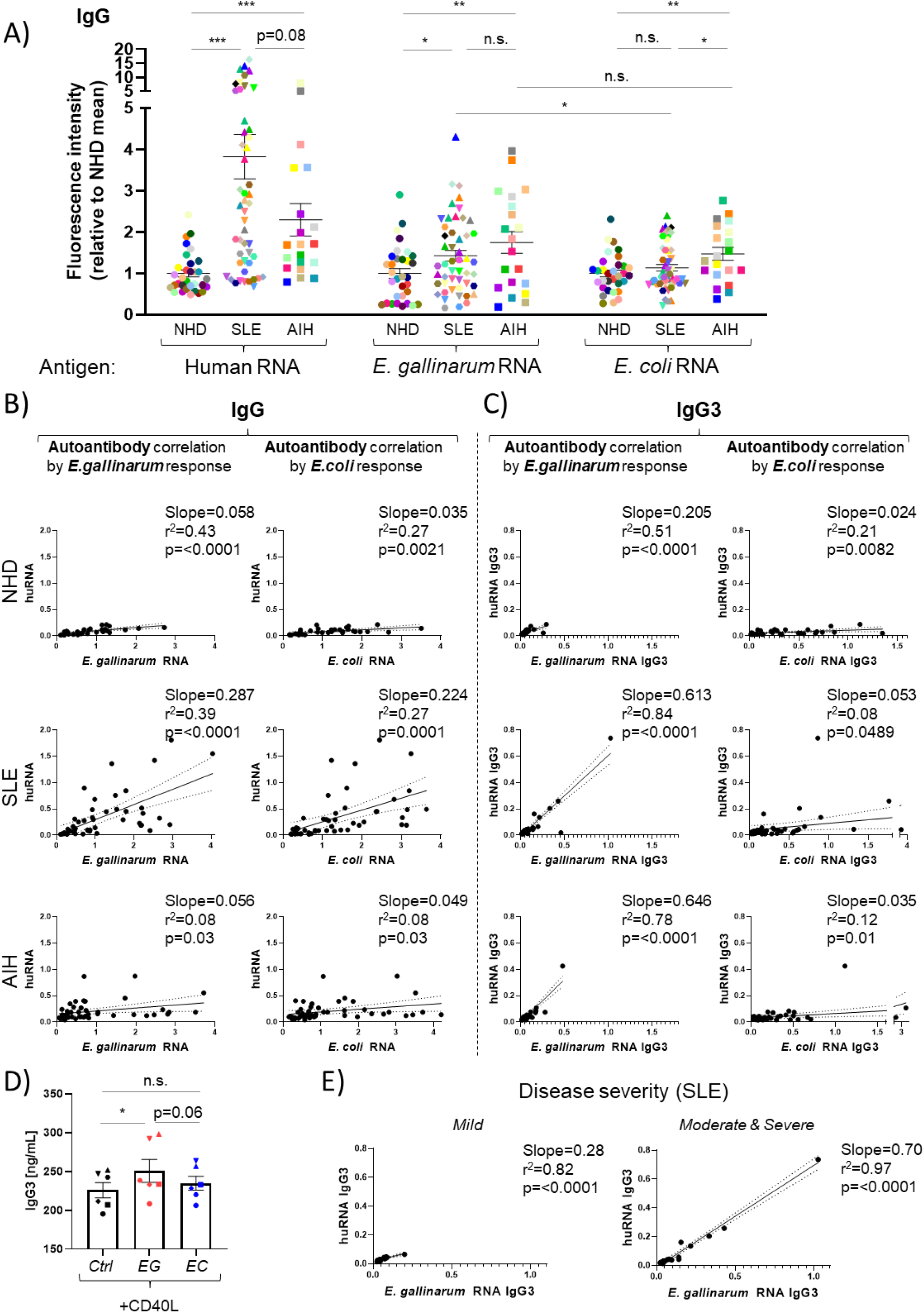
SLE and AIH patients mount IgG and IgG3 responses against *E. gallinarum* RNA. Plates were coated with RNA preparations from human PBMCs (Human RNA), *E. gallinarum* (*E. gallinarum* RNA) or *E. coli* (*E. coli* RNA) bacterial cultures. The reactivity of human serum was measured by ELISA. a) Total IgG reactivity of sera from normal healthy donors (NHD), systemic lupus erythematosus patients (SLE) and autoimmune hepatitis patients (AIH) was tested against different RNA preparations as indicated (NHD n=32, SLE n=49, AIH n=20). Results were normalized to the average of NHD controls for each RNA preparation. b) Correlation between absolute reactivity of total IgG against human RNA and *E. gallinarum* RNA (left panels) or *E. coli* RNA (right panels) for each group (NHD, SLE, AIH). c) Correlation between absolute reactivity of IgG3 against human RNA and *E. gallinarum* RNA (left panels) or *E. coli* RNA (right panels) for each group (NHD, SLE, AIH). d) Human PBMCs were stimulated with *E. gallinarum (EG)* or *E. coli (EC)* lysates in the presence of anti-CD3/CD28 and CD40L activation. Supernatants were taken on day 7 and IgG3 concentrations were measured by ELISA. e) Correlation between absolute reactivity of IgG3 against human RNA and *E. gallinarum* RNA for SLE patients divided based upon their disease severity (Mild, Moderate/Severe). Statistical analysis: Unpaired t-test (a); paired t-test (d); linear regression (b,c,e)

To assess whether anti-gut bacterial RNA responses were heightened more generally, we evaluated the relative reactivity against *E. coli* RNA, which is commonly found in intestinal microbiomes. There was significantly less reactivity towards *E. coli* RNA compared to *E. gallinarum* RNA in SLE patient sera (Fig. 7A). Also, reactivities to *E. coli* RNA were not different in normal healthy donors (NHD) and SLE patients but higher in AIH, suggesting preferential targeting of *E. gallinarum* in SLE compared to AIH (Fig. 7A).

To test whether the increased reactivities to *E. gallinarum* RNA were due to higher titers of total IgG in autoimmune patients (fig. S7A), we correlated the absolute IgG titers with reactivity against the tested antigens. There was no correlation in any patient group (fig. S7B), supporting a focused IgG response to this pathobiont.

To differentiate between signs of chronic immune stimulation and intermittent acute responses against translocating bacteria, we assessed IgM levels towards the gut bacteria. IgM levels were elevated against *E. gallinarum* as well as *E. coli* in SLE patients (fig. S7C). The strong correlation between those reactivities suggests constantly ongoing immune responses against these bacteria in the gut, possibly as a result of an impaired intestinal barrier (fig. S7D) (*18, 63, 64*).

When assessing whether chronic systemic immune responses with IgG class switching reveal more selective associations, we found tightly correlated IgG responses between anti-*E. gallinarum* IgG and anti-human RNA IgG autoantibodies in SLE but not AIH patients with negligible antibody titers in healthy subjects (Fig. 7B).

While there was no association between disease severity and autoantibodies, we detected increased IgG titers against *E. gallinarum* and *E. coli* RNA during SLE flares (fig. S7, E and F). Because reactivities to both bacteria increased, we suspect that this resulted from a deteriorating intestinal barrier during flares given that an increase in inflammatory stimuli impairs gut barrier function (*3*).

In the lupus-prone murine models, the IgG3 subclass robustly correlated with disease activity and proteinuria *in vivo* (see Fig. 6I). While IgG subclasses differ between humans and mice (*65*), IgG3 can be induced by IL-17 in humans (*66*) and is most potent for FcR and complement activation (*67, 68, 69*), which are both critical in the pathophysiology of SLE.

When we tested the contribution of different IgG subclasses to anti-RNA reactivity, we found a trend towards higher IgG3 autoantibodies as well as anti-*E. gallinarum* IgG3 titers in SLE patients (fig. S7G). In addition, it was notable that individual subjects exhibited high reactivities against *E. gallinarum*-RNA as well as self-RNA. Furthermore, we correlated those reactivities for all patient groups and detected a tight correlation between IgG3 responses against human RNA and *E. gallinarum* RNA (Fig. 7C). In contrast, anti-*E. coli* RNA IgG3 did not associate with autoantibodies in SLE or AIH patients, emphasizing the connection between anti-*E. gallinarum* IgG3 reactivity and autoantibody load. This finding was unique for IgG3 as it could not be observed for any other IgG subclass (fig. S7H). The high IgG3 reactivity against bacterial- and self-antigens were also not dependent on total IgG content in the serum (fig. S7I).

Finally, we tested whether the induction of IgG3 by *E. gallinarum* can also occur in human immune cells. To this end, we measured IgG3 levels in the supernatants of PBMC from healthy subjects that were stimulated with *E. gallinarum* in the presence of CD40L. After 7 days, there was a small but significant increase in IgG3 in the supernatants of *E. gallinarum*-stimulated cultures compared to controls (Fig. 7D). We also saw a trend towards higher IgG3 content compared to *E. coli*-stimulated cultures, despite those cultures containing massively more mature B cells after CD40L costimulation (see Fig. 5E).

To determine if higher IgG3 autoantibody and anti-*E. gallinarum* titers were connected to clinical parameters, we separated the SLE patients according to their disease severity. All patients with high anti-*E. gallinarum* IgG3 and self-RNA IgG3 belonged to the group with moderate and severe disease severity (Fig. 7E). These results indicated that IgG3 responses are associated clinically with more severe disease in SLE patients. Taken together, the patient-derived data demonstrates a strong correlation between antibodies targeting autoantigens and responses against *E. gallinarum*; this connection was particularly prominent for antibodies of the IgG3 subclass, which could be induced in human PBMC by *E. gallinarum*.

## Discussion

Gut pathobionts contributing to immune-mediated diseases are increasingly identified in animal models or implicated based on human association studies (reviewed in (*3, 70*). Progress has yet to be made, however, in elucidating the effects of gut pathobionts on human immune responses. Human host-microbiota interactions are little studied except for cross-reactivity (*17, 56, 71, 72*). We have shown that the translocating gut pathobiont *E. gallinarum* promotes human Th17 differentiation *in vitro* by inducing Th17 differentiation factors in human monocytes via a contact-dependent mechanism. This effect is inhibited by endosomal TLR blockade, specifically by TLR8 inhibition and the drug HCQ that is commonly used in lupus patients and known to dampen endosomal TLR signaling (*49*). In addition, IRAK4 inhibition abrogated Th17 induction, which is a key signaling molecule downstream of TLRs with inhibitors being in development for SLE (*73*). IRAK4 promotes endosomal TLR signaling in human APCs and deficiency of IRAK4 in mice attenuates lupus-like disease models (*74, 75*). Human TLR7/8, upstream of IRAK4, are highly implicated in SLE and scleroderma pathogenesis (*76, 77*), but the triggers remain ill-defined. Among endogenous and viral RNA, bacterial RNA structures are plausible candidates (*51, 78*). Interestingly, TLR8 senses microbial viability which triggers T follicular helper (Tfh) differentiation and systemic antibody responses (*51*). Also, TLR8 reverses regulatory T cell function (*79*), which could add further to the breach in immunologic tolerance in systemic autoimmunity. The role of *E. gallinarum* RNA in TLR7/8 triggering warrants further study, given that anti-*E. gallinarum* RNA IgG responses are induced in SLE and AIH patients, and are highly correlated with anti-RNA autoantibody titers. The contact-dependent, non-soluble nature of *E. gallinarum*-induced monocyte activation and Th17 differentiation suggests that RNA or related molecular structures in special bacterial compartments may be responsible for this phenomenon (*80*).

For any *E. gallinarum*-related molecular structures to trigger endosomal TLRs, it is expected that *E. gallinarum* is taken up by monocytes but escapes rapid elimination as supported by host immune evasive mechanisms recently demonstrated (*19*). Mucosally adapted *E. gallinarum* strains that evade immune destruction evolve within the host to translocate to secondary lymphoid organs and the liver. In tissues and blood, they may promote maladaptive Th17 responses that contribute to extraintestinal autoimmunity. Here, we show that multiple *E. gallinarum* strains from different host species promote human Th17 differentiation. It is therefore likely that the maladaptive Th17 responses *in vivo* are broadly mediated by any *E. gallinarum* strain once translocated after within-host evolution in the small intestine (*19*).

Th17 cells can have pro- or anti-inflammatory effects depending on IL-10 or IFNγ production (*2, 37, 81, 82, 83, 84, 85*). We have shown that *E. gallinarum*-induced human Th17 cells co-produce IFNγ but downregulate IL-10, supporting a proinflammatory phenotype. This functional phenotype was induced in healthy human cells by *E. gallinarum* compared to other bacteria that only skew this balance during an inflammatory state (*83*). IL-1β is one of the factors regulating the balance between these “flavors” of Th17 cells (*2, 86, 87*), which was also induced by *E. gallinarum* co-cultured with human monocytes.

Once pro-inflammatory Th17 cells are induced, autoreactive germinal centers can develop and support Tfh and autoantibody production in susceptible hosts (*25, 26*). Th17 cells preferentially drive B cells to switch to IgG2a and IgG3 subclasses (*57*). We found that *E. gallinarum* induces particularly autoantibodies of the IgG3 subclass with tight correlations between anti-human RNA autoantibodies and anti-*E. gallinarum* RNA in sera of SLE and AIH patients, respectively. We demonstrated that systemic IgG3 autoantibody responses in gnotobiotic mouse models are linked to autoimmune kidney damage and proteinuria *in vivo*. IgG3 is a significant subclass of several autoantibodies in connective tissue diseases (*88, 89, 90*) and autoimmune liver diseases (*91, 92, 93*). Future work is needed to determine if the progressive translocation of *E. gallinarum* to secondary lymphoid organs and the liver triggers hepatic and systemic IgG3 immune complexes depositing in the kidneys that lead to glomerulonephritis. A systemic maladaptive IgG3 response to *E. gallinarum* and similar translocating pathobionts is likely, given the known effector functions triggered by bacterial infections, including complement activation and cytotoxicity (reviewed in (*67*)).

Our study was limited by a focus on human Th17 responses from healthy subjects. Future work should determine if *E. gallinarum* induces similar (or even more pronounced) proinflammatory responses in autoimmune patients and if single cell analyses reveal heterogeneity within bulk Th17 cells. The human autoantibody profiles linking *E. gallinarum* responses *ex vivo* were patient-derived, suggesting similar mechanisms in autoimmune patients, but will need to be confirmed in larger cohort studies, including analyses of potential confounding factors such as microbiome composition, environmental influences and genetic predispositions. *E. gallinarum* prevalence in the stool of patient cohorts also needs to be determined despite the caveat that translocating pathobionts do not necessarily shed easily in the stool once mucosally adapted to its niche (*18, 19*). Lastly, the molecular mechanisms need to be dissected in more depth with genetic manipulation of the TLR8 pathway and identification of the microbial structures within *E. gallinarum* that skew T helper cell differentiation in a monocyte contact-dependent manner. Despite the need for more molecular insights, the cellular mechanisms we uncovered in this translational study support a conceptual framework in autoimmunity that links *E. gallinarum*-mediated monocyte-Th17 skewing and IgG3 autoantibody production in humans (Fig. 7J).

We focused here on the translocating pathobiont *E. gallinarum* but other gut commensal bacteria such as *Lactobacillus reuteri* have also been shown to translocate to lymphoid tissues and the liver in autoimmune-prone hosts (*55, 94*). Future studies should dissect additional human immune responses induced by these and other translocating pathobionts that are expected to directly interact with host immune cells. These efforts may eventually lead to a better understanding of human host-microbiota interactions, which may inform new drug development on the host side (*49*) as well as new targets in the human microbiome that could be removed by precision approaches (*95, 96*).

## Acknowledgments

We are grateful to V. Hoelt, P. Vallet, V. Haas, T. Hoenig, and A. Schneider for excellent technical support. The authors would also like to thank A. Zipperer, P. Scepanovic and A. Mechling for insightful discussions on this topic. We also acknowledge the support of the Roche animal facility and Yale Animal Resources Center (YARC) for expert animal care. M.N. was supported by a grant from the National Institutes of Health (NIH NIAID F30AI157227). Part of the work was supported by grants from the Lupus Research Alliance (Novel Research Grant and Lupus Insight Prize), the National Institutes of Health (NIH R01AI118855, T32AI07019), Arthritis National Research Foundation, Arthritis Foundation, and the Maren Foundation (all to M.A.K.).

## Author Contributions

K.G., N.P., C.B. and M.A.K. conceptualized the project and methodology. K.G., M.N., N.Sa., J.S., Y.Y., N.So., S.L., A.L.M., R.H., D.D., M.F., Do.S. and S.M.V. performed experiments and analyses. K.R., G.M.D., Da.S., C.S., K.G.L., L.P. participated in the conceptualization of the project, provided access to resources, clinical samples and critical intellectual input. K.G., L.P., C.B., and M.A.K. wrote the manuscript with input from all authors.

## Materials & Methods

### Bacterial culture

Bacteria were plated on Mueller-Hinton agar plates and colonies were picked to be grown in Gifu liquid medium overnight at 37°C. Cultures were pelleted and washed three times with PBS. The optical density at 600 nm (OD_600_) was measured for calculations of colony forming units (CFUs). Bacterial solutions were sonicated three times with Branson Digital Sonifier at 60% for 30 seconds. Complete inactivation of bacteria was controlled by plating and ensured by adding penicillin/streptomycin to eukaryotic cultures. These bacterial lysates were added to PBMC cultures equivalent to the PBMC:CFU ratio of 1:1.

For heat inactivation, bacterial lysates were denatured at 80°C for 40 minutes.

To separate bacterial culture lysate and supernatant, bacteria were grown in RPMI 1640 medium overnight. Supernatants were filtered (0.2 μM), sonicated and subsequently added to PBMC cultures. Corresponding bacteria were lysed as described above.

For treatment with lysozyme (1 mg/mL, Alfa Aesar) or proteinase K (Roche), bacteria were resuspended in the provided buffer before sonication. After sonication, enzymes were added and incubated at 37°C for 1 hour for lysozyme and at 55°C for 1 hour for proteinase K. The reaction was stopped by heating to 65°C for 20 minutes for lysozyme and to 95°C for 10 minutes for proteinase K. Sodium periodate was dissolved in sodium acetate buffer. Bacterial lysates were incubated with 1 mM sodium periodate overnight protected from light and subsequently purified using Zeba Spin DeSalting columns (ThermoFisher).

### Sources of bacterial isolates and strains

| Bacteria | Source |
| --- | --- |
| ***Enterococcus gallinarum* isolates** |  |
| *E. gallinarum* isolate from NZW x BXSB F<sub>1</sub> hybrid mice | Manfredo Vieira et al. (18) |
| DSMZ 20718 | German Collection of Microorganisms and Cell Cultures GmbH (DSMZ) |
| DSMZ 25780 |  |
| DSMZ 28564 |  |
| DSMZ 100110 |  |
| ATCC 49573 | American Type Culture Collection (ATCC) |
| ATCC 49609 |  |
| ATCC 700425 |  |
| E10_172_EG_BA4 | National Collection of Industrial, Food and Marine Bacteria |
| E20_190_EF_BA4 |  |
| ***Enterococcus faecalis* isolate** |  |
| ATCC 51299 | American Type Culture Collection (ATCC) |
| ***Enterococcus casseliflavus* isolate** |  |
| ATCC700668 | American Type Culture Collection (ATCC) |
| ***Escherichia coli* isolate** |  |
| ATCC25922 | American Type Culture Collection (ATCC) |

### PBMC cultures

Blood was drawn from healthy human subjects and collected in lithium heparin tubes (Becton Dickinson). PBMCs were isolated using Ficoll-Paque PLUS (Cytiva). For proliferation analysis, cells were incubated with the CellTrace Violet Cell Proliferation Kit (ThermoFisher). PBMCs were resuspended in RPMI 1640 medium with 4.5 g/L D-glucose, 2.383 g/L HEPES buffer, L-glutamine, 1.5 g/L sodium bicarbonate, 110 mg/L sodium pyruvate (ThermoFisher) supplemented with 10% FCS, 2% human serum (Sigma-Aldrich), 1% non-essential amino acids (ThermoFisher), 100 Units/mL penicillin, 100 ug/mL streptomycin (Gibco). Either 5U IL-2 (Peprotech) or ImmunoCult Human CD3/CD28 T cell activator (Stemcell Technologies) was added at a dilution of 1:500.

Blocking antibodies, inhibitors, and cytokines were added to the cultures (concentrations in table below). Bacterial lysates were prepared as described above and added to the cultures. Cells were incubated at 37°C, 5% CO_2_ for seven days. For intracellular cytokine analyses, cells were stimulated with PMA/ionomycin with Brefeldin A (1:500 or equivalent to 1.3 μM ionomycin and 81 nM PMA, Cell Activation Cocktail, Biolegend) for 4 hours. Cells were washed and stained for viability (Fixable viability dye eFlour 780, eBioscience), afterward kept in staining buffer (2% FCS, BD Biosciences). Surface proteins were stained for 30 minutes at 4°C, cells were subsequently fixed in the FoxP3/Transcription factor staining buffer set (eBioscience) overnight. Intracellular proteins were stained for 90 minutes at 4°C. Cells were analyzed with a BD Biosciences Fortessa cell analyzer. Plasmacytoid DCs (pDCs) were defined as CD3^-^CD19^-^ CD11c^Med^CD123^+^MHCII^+^, CD14^+^ monocytes as CD3^-^CD19^-^CD123^-^CD16^-^CD14^+^, CD16^+^ monocytes as CD3^-^CD19^-^CD123^-^CD16^+^CD14^-^, classical DCs (cDCs) as CD3^-^CD19^-^CD123^-^ CD11c^+^MHCII^+^ and B cells as CD3^-^CD19^+^. Human B cell populations were defined as naïve B cells CD19^+^CD27^-^CD38^-^, memory B cell CD19^+^CD27^+^CD38^-^, plasmablasts CD19^+^CD27^+^CD38^+^ and plasma cells CD3^-^CD19^low^CD138^+^.

Secreted proteins in the supernatant were measured with a custom ProcartaPlex multiplex immunoassay kit on a Luminex analyzer according to the manufacturer’s instructions (both ThermoFisher).

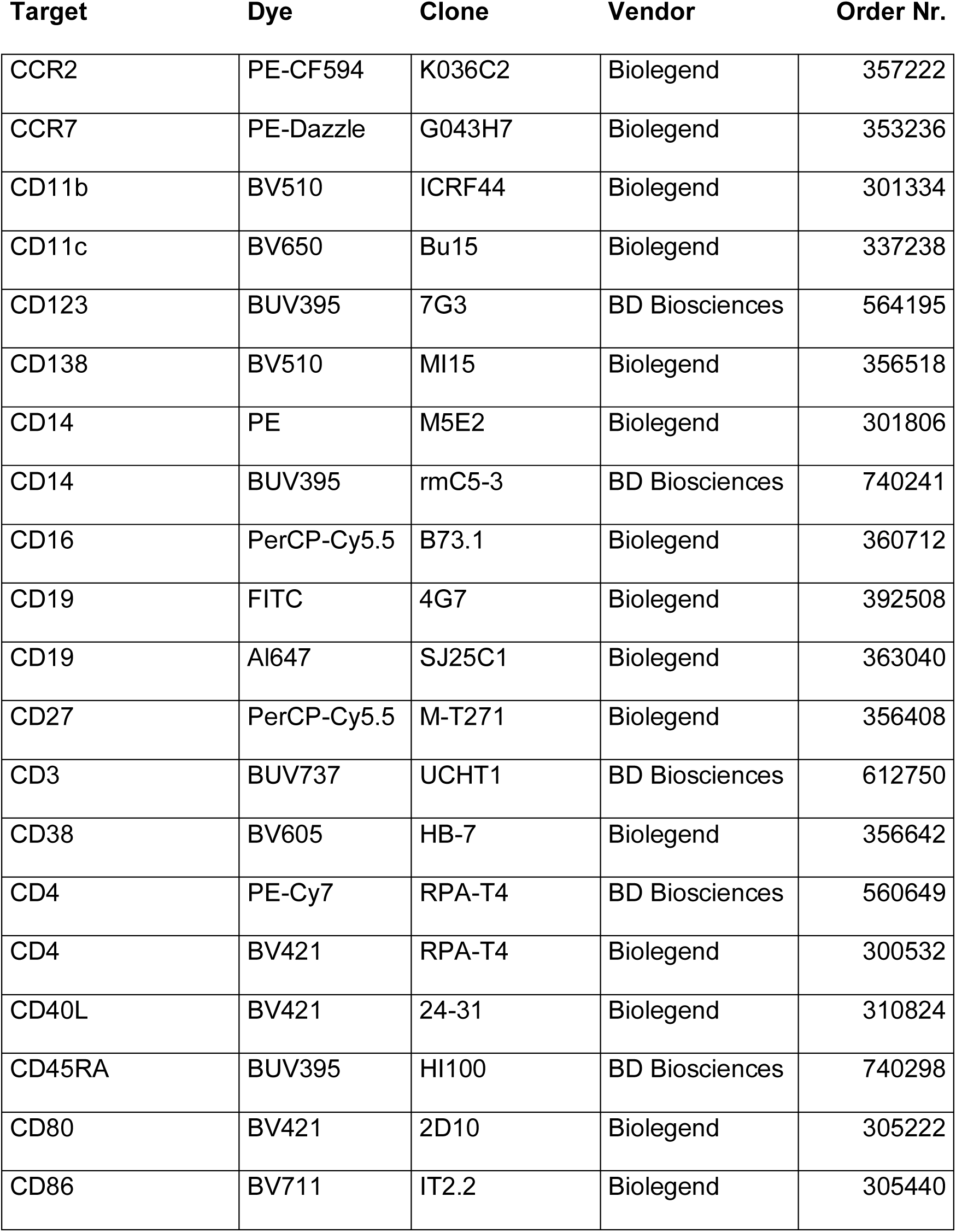
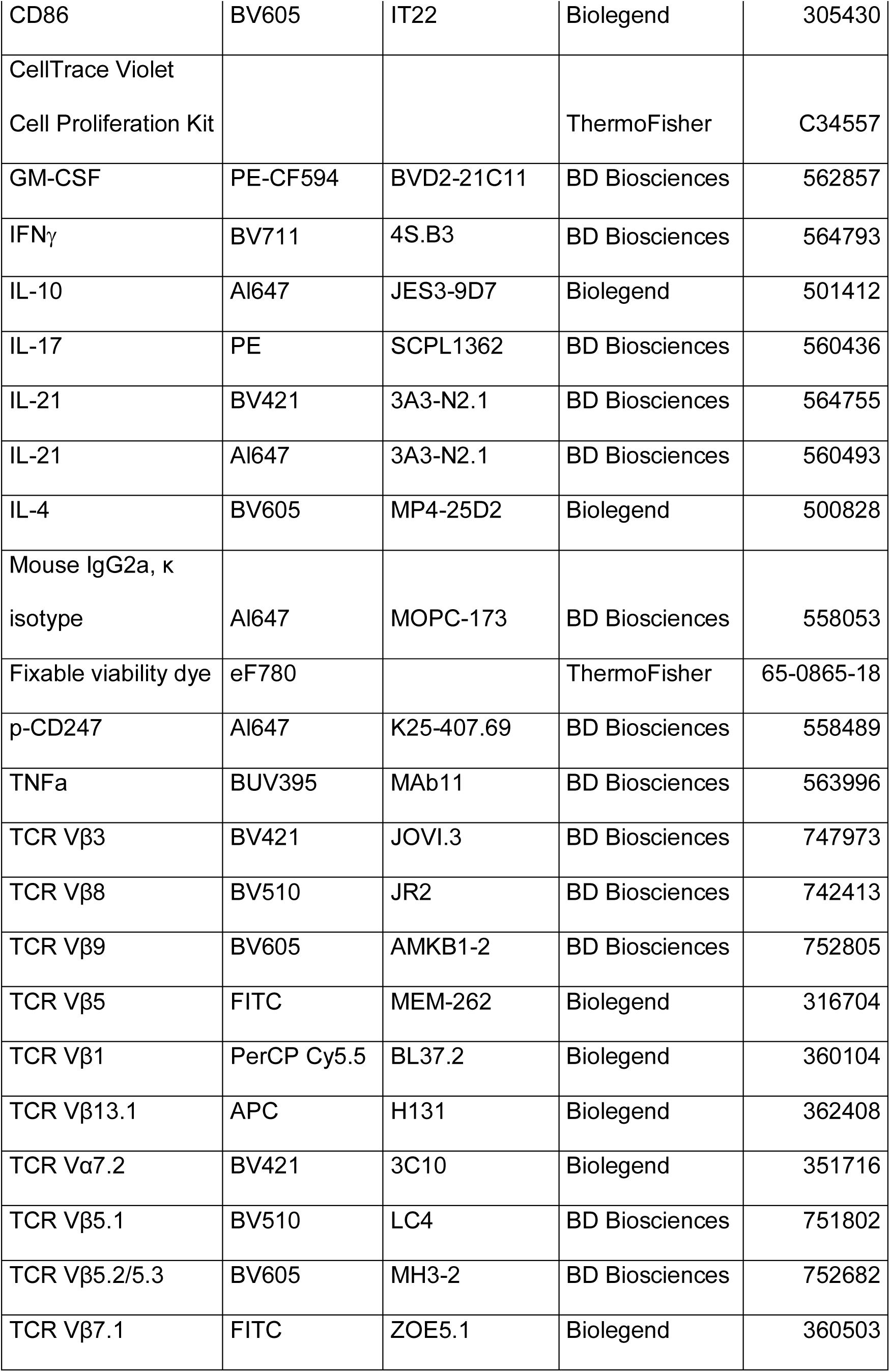
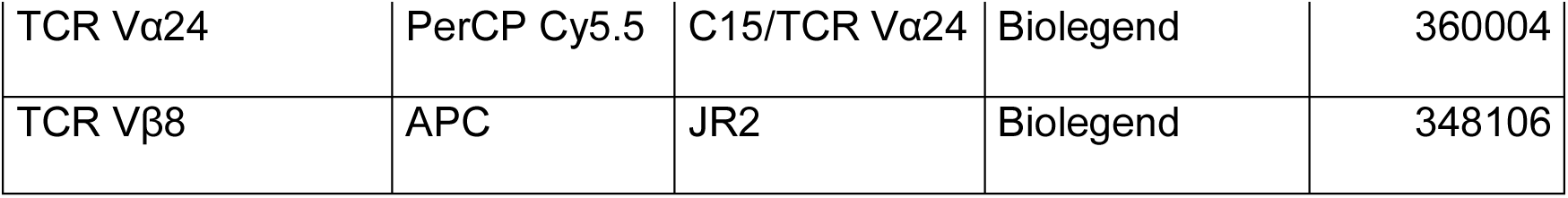

### Stimulants and inhibitors

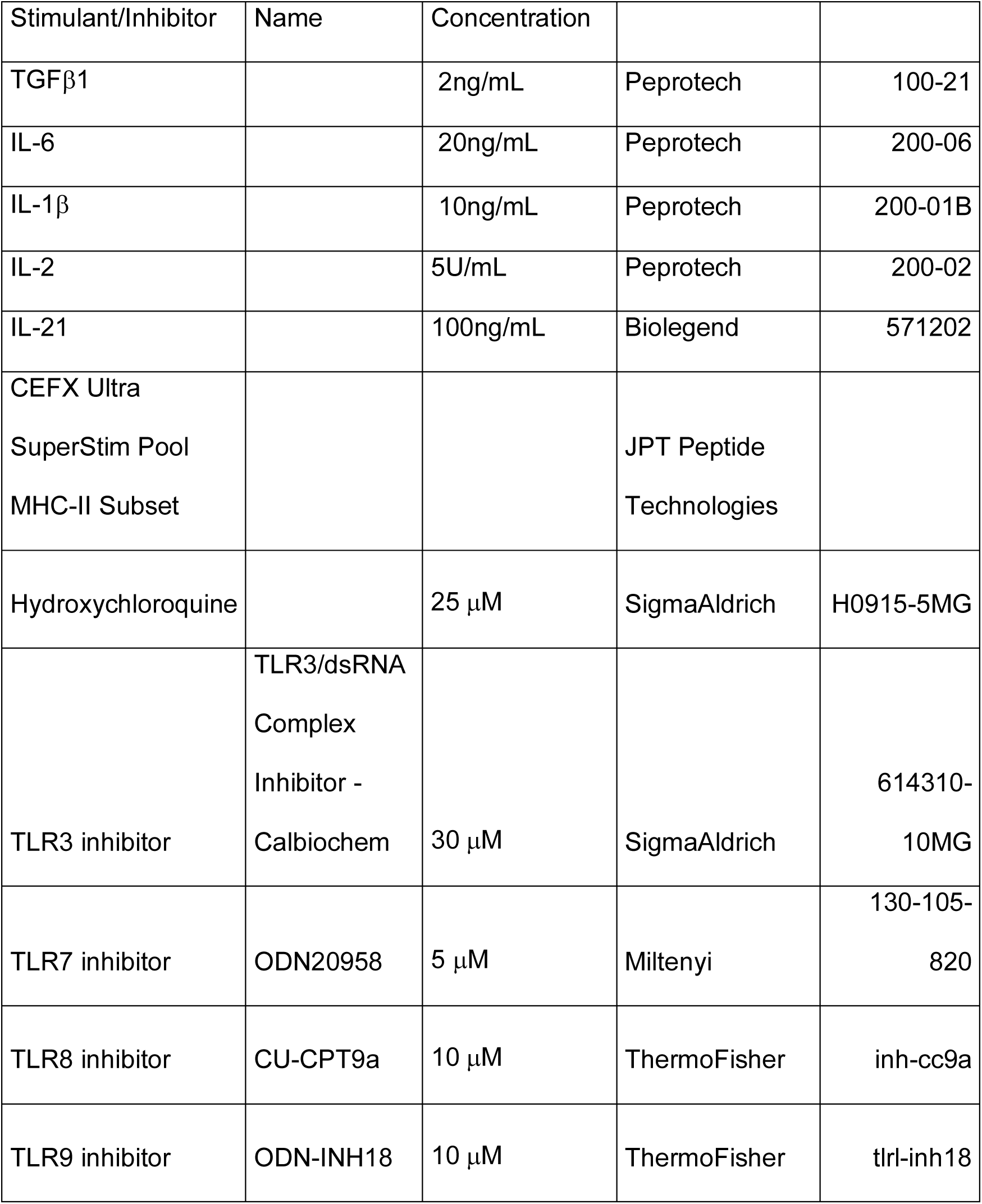
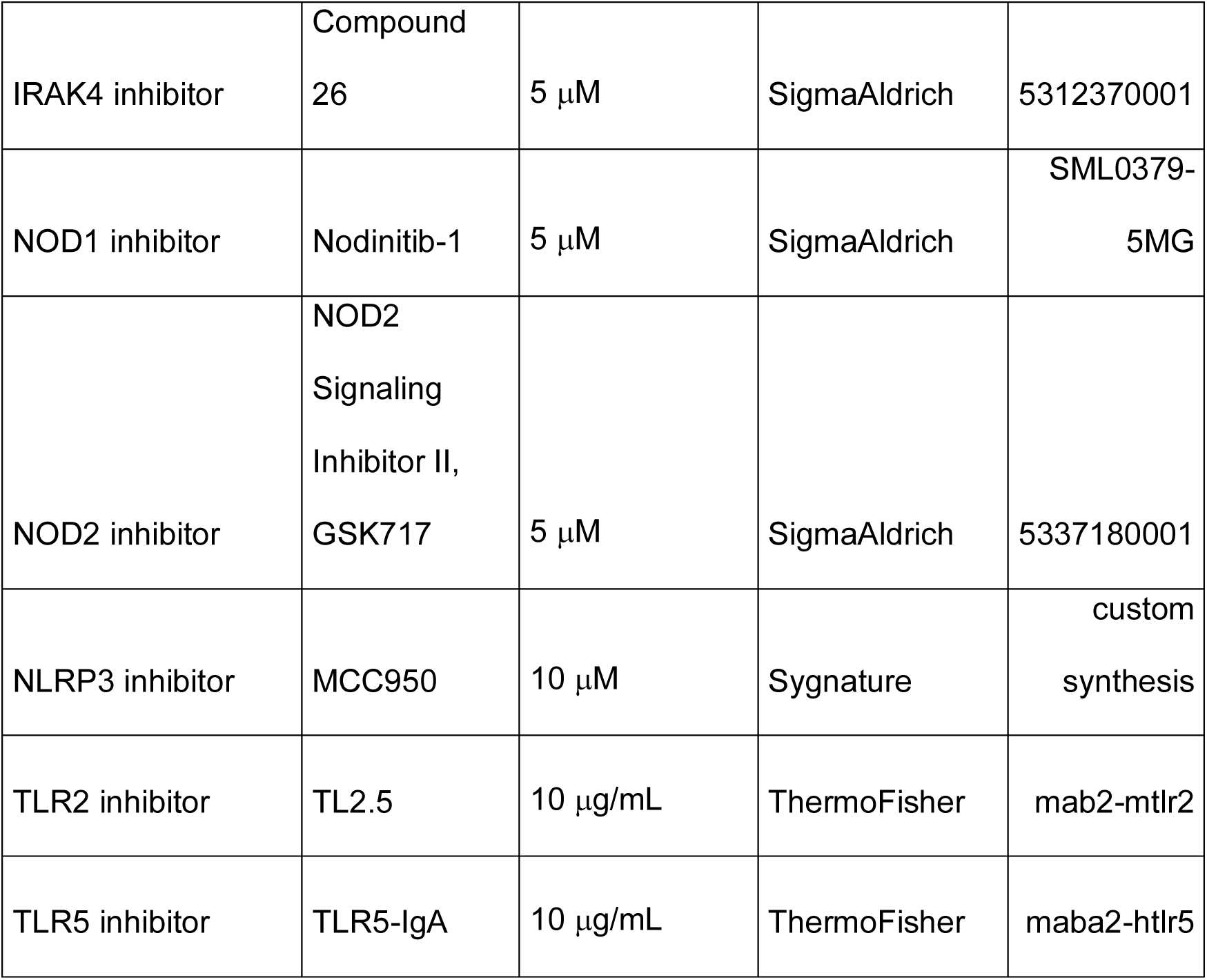

| Stimulant/Inhibitor | Name | Concentration |  |  |
| --- | --- | --- | --- | --- |
| TGFβ1 | | 2ng/mL | Peprotech | 100-21 |
| IL-6 |  | 20ng/mL | Peprotech | 200-06 |
| IL-1β | | 10ng/mL | Peprotech | 200-01B |
| IL-2 |  | 5U/mL | Peprotech | 200-02 |
| IL-21 |  | 100ng/mL | Biolegend | 571202 |
| CEFX Ultra<br>SuperStim Pool<br>MHC-II Subset |  |  | JPT Peptide<br>Technologies |  |
| Hydroxychloroquine | | 25 μM | SigmaAldrich | H0915-5MG |
| TLR3 inhibitor | TLR3/dsRNA<br>Complex<br>Inhibitor -<br>Calbiochem | 30 μM | SigmaAldrich | 614310-10MG |
| TLR7 inhibitor | ODN20958 | 5 μM | Miltenyi | 130-105-820 |
| TLR8 inhibitor | CU-CPT9a | 10 μM | ThermoFisher | inh-cc9a |
| TLR9 inhibitor | ODN-INH18 | 10 μM | ThermoFisher | tlrl-inh18 |
| IRAK4 inhibitor | Compound<br>26 | 5 μM | SigmaAldrich | 5312370001 |
| NOD1 inhibitor | Nodinitib-1 | 5 μM | SigmaAldrich | SML0379-5MG |
| NOD2 inhibitor | NOD2<br>Signaling<br>Inhibitor II,<br>GSK717 | 5 μM | SigmaAldrich | 5337180001 |
| NLRP3 inhibitor | MCC950 | 10 μM | Sygnature | custom<br>synthesis |
| TLR2 inhibitor | TL2.5 | 10 μg/mL | ThermoFisher | mab2-mtlr2 |
| TLR5 inhibitor | TLR5-IgA | 10 μg/mL | ThermoFisher | maba2-htlr5 |

### B cell cultures

PBMCs were isolated as described above. For B cell cultures, CD40L-trimer (2 nM, EnzoLifeScience) and IL-21 (100 ng/mL, Peprotech) were added. Supernatant was taken on day 7 and stored at −80°C until analysis. B cell populations were analyzed by flow cytometry as described above. IgG3 concentrations were measured by an IgG3 Human ELISA Kit (Invivogen).

### Isolation of cell populations

PBMCs were isolated as described above. Cells were divided into two fractions for magnetic cell separation. CD4^+^ T cells were isolated from one fraction with the EasySep Human CD4^+^ T Cell Isolation kit, and CD14^+^CD16^-^ monocytes were isolated from the other fraction using the EasySep Human Monocyte Isolation kit. Separation was carried out with a RoboSep-S according to the manufacturer’s instructions (all Stemcell Technologies).

For co-culture experiments, 200,000 CD4^+^ T cells were combined with 20,000 or 100,000 CD14^+^ monocytes. For transwell cultures (0.3 μM pore size), CD14^+^ monocytes were added to the lower well, while CD4^+^ T cells were seeded onto the upper inlet. The bacterial lysate was added to both chambers.

### Epithelial barrier integrity studies

HT29-MTX (*97*) cells were cultured in DMEM-F12 GlutaMax with 10% FCS and 1% non-essential amino acids. Caco-2 (ATCC) cells were cultured in DMEM-F12 GlutaMax with 20% FCS and 1% non-essential amino acids. 10,000 epithelial cells were seeded per well into an xCELLigence 16-well E-plate (well diameter 10.65 mm). Cell index, as indicated by impedance, was monitored real-time using the xCELLigence Real-Time Cell Analyser. Cells were grown until the cell index reached stability. Medium was replaced every 48 hours and additionally supplemented with 100 mM HEPES buffer. Living bacteria were resuspended in 20 μL DMEM-F12 GlutaMax per well and added to the epithelial cultures. Impedance was measured every 30 minutes for at least 24 hours.

### ELISA

RNA was extracted from *E. gallinarum* and *E. coli* cultures with AllPrep PowerFecal DNA/RNA Kit (QIAGEN) according to the manufacturer’s instructions. RNA from human PBMCs was isolated using RNeasy Kit (QIAGEN) according to the manufacturer’s instructions. RNA was plated at 500 ng/well (10 ng/μL) on Thermo Scientific CovaLink and Immobilizer Amino plates (ThermoFisher) overnight at 4°C. Loading was normalized by using an anti-POLR2A antibody (R&D Systems) to test reactivity to coated RNA.

For murine serum analysis, salmon sperm DNA was used for dsDNA and Yeast RNA (both ThermoFisher) was used for RNA antigen coating. Serum was collected and stored at −20°C until further use. Anti-dsDNA and anti-RNA autoantibodies in serum were assessed by ELISA as previously described (*18*). Briefly, plates were washed and blocked for 90 minutes with PBS containing 0.05% Tween-20, 5% FCS and 1% BSA. Plates were washed again and incubated for two hours with the respective serum, diluted 1:1000 in PBS with 1% BSA. Finally, plates were washed again and incubated with secondary detection antibody conjugated to HRP in PBS with 1% BSA for 1 hour. Serum reactivity was measured with TMB substrate (ThermoFisher) on a Tecan Spark microplate reader (Tecan).

IgG concentration in serum was measured with human IgG ELISA Kit (Abcam). In addition, IgG3 concentration in culture supernatants was measured with an ELISA Kit for IgG Subclasses (ThermoFisher).

### Antibody sources

| **Target** | **Dilution** | **CatNr.** | **Vendor** |
| --- | --- | --- | --- |
| pan-IgG | 1:2000 | 62-8420 | Thermo |
| pan-IgA | 1:2000 | PA1-74395 | Thermo |
| IgM | 1:500 | 05-4920 | Thermo |
| IgG1 | 1:1000 | MH1715 | Thermo |
| IgG2 | 1:1000 | MH1722 | Thermo |
| IgG3 | 1:4000 | MA5-16718 | Thermo |
| IgG4 | 1:1000 | MH1742 | Thermo |
| Mouse<br>IgG | 1:500 | G-21040 | Thermo |
| Anti-<br>sheep | 1:500 | A16041 | Thermo |
| dsDNA | 1:1000-32,000 | MAB1293 | EMD Milipore |

### Human healthy donor and patient characteristics*

Human healthy donor and patient characteristics\*
|  | Healthy<br>Donor Pool 1<br>(Basel) | Healthy<br>Donor Pool 2<br>(Yale) | SLE (Chemie<br>Brunschwig) | SLE (Yale) | AIH<br>(Hamburg) |
| --- | --- | --- | --- | --- | --- |
| Age | 42.02±11.32 | 38±12 | 53±12 | 42.5±12.1 | 55.83±15 |
| Sex | 50.25% Male<br>49.25%<br>Female | 0% Male<br>100%<br>Female | 12% Male<br>88% Female | 0% Male<br>100%<br>Female | 33% Male<br>66% Female |
| Ethnicity | n/a | 45% White<br>45%<br>Hispanic | 87% White<br>10% Hispanic<br>3% Other | 31% White<br>25%<br>Hispanic | 100% White |
|  |  | 10% Other |  | 44% Black |  |
| Anti-dsDNA | n/a | n/a | 100% | 50% | n/a |
| Disease<br>Severity | n/a | n/a | 45% Mild<br>48%<br>Moderate<br>7% Severe | n/a | n/a |
| Disease<br>Flare | n/a | n/a | 44% | 19% | 50% acute<br>disease |

\*n/a, not assigned.

Disease severity was determined by the physician’s global clinical assessment (PGA) at the time of appointment (*98*). The healthy donor pool 2 (Yale) and the SLE patient cohort (Yale) were partly matched. Both SLE patient cohorts, SLE (Chemie Brunschwig) and SLE (Yale) were not related and represent independent patient populations. The healthy donor pool 1 and the AIH cohort were also independent from the other cohorts. Results have been confirmed in both SLE cohorts independently and data is shown as pooled analyses of all cohorts. All human subject protocols were conducted in accordance with the principles of the Declaration of Helsinki and approved by the respective institutional ethics committee at each center. A signed document of informed consent was obtained from all study participants.

### Monocolonization experiments

Germ-free C56BL/6 female mice were maintained in flexible film isolators and given irradiated and autoclaved standard chow diet (Teklad 2018S) as recently described (*19*). Mice were on a dark/light cycle of 12 hours/12 hours with controlled temperature and humidity. Monocolonization experiments were conducted by transferring germ-free mice from flexible film isolators to positive pressure ventilated microisolator cages (Techniplast #ISO72P). Each mouse was immediately inoculated with approximately 6 × 10^7^ CFU bacterium of interest upon transfer. Bacterial inocula were freshly prepared the day of gavage by subculturing bacteria and grown for 4 hours at 37°C. All animal protocols were approved by the Yale University Institutional Animal Care and Use Committee (IACUC Protocol 2018-11513). Immediately after gavage, topical treatment of 1.25 mg 5% imiquimod (IMQ) cream was administered on the skin of the ear (alternating between left and right ears) every other day for 8 weeks. Animal care and handling was approved by the Yale Institutional Animal Care and Use Facility and in accordance with the National Institutes of Health (NIH) Guide for the Care and Use of Laboratory Animals.

### Organ weights, bacterial translocation and histological scoring

After 8-weeks of monocolonization and imiquimod treatment, serum was collected under isoflurane anesthesia by retro-orbital puncture. Mice were then euthanized and the liver, mesenteric lymph nodes, and spleen were aseptically dissected and weighed. Liver, mesenteric lymph node, and spleen homogenates were generated by mechanical digestion through a 70-μM mesh filter. Aliquots of homogenates were reserved to assess bacterial translocation by plating on Gifu agar plates and cultured anaerobically for 48 h. *E. gallinarum* isolates were confirmed by species-specific PCR as recently described (*19*).

For histology scoring, the following method was used: Slides were examined by a board-certified veterinary pathologist. Livers were evaluated for inflammation, necrosis, and fibrosis using the revised Knodell scoring system (a worksheet is found at https://tpis.upmc.com/tpislibrary/schema/mHAI.html). Kidneys were evaluated for glomerulonephritis and assigned a severity score of 1 to 4 in which 1=minimal, 2=mild, 3=moderate, 4=severe. Other microscopic findings were identified and assigned a severity of minimal, mild, moderate, or severe.

### Flow cytometry of murine cells

Lymphocytes were isolated from the mesenteric lymph nodes, spleen, lamina propria, and peyer’s patches as previously described (*18*) (*99*). Isolated lymphocytes were stimulated with PMA/ionomycin with Brefeldin A (1:500 or equivalent to 1.3 μM Ionomycin and 81 nM PMA, Cell Activation Cocktail, Biolegend) for 5 h at 37°C with 5% CO_2_. Cells were surface stained in a 1% BSA solution for 20 min at 4°C. For intracellular staining, cells were fixed in BD Cytofix/Cytoperm kit (BD Biosciences) for 10 minutes and intracellular proteins were subsequently stained overnight at 4°C. Cells were analyzed with a BD CytoFLEX cell analyzer. Murine B cell populations in the spleen were defined according to (*100*) as CD19^int^CD138^+^B220^int^ plasmablasts, CD19^int^CD138^+^B220^low^ early plasma cells and CD19^low^CD138^+^B220^low^ mature plasma cells.

### Specific-pathogen free (SPF) mouse experiments

SPF C56Bl/6 female mice were obtained from Charles River laboratories at the age of 6 weeks. Mice were kept in individually ventilated cages, acclimated for one week, and microchipped for identification. Mice were treated for two weeks with oral antibiotics (vancomycin (0.5 g/L, SigmaAldrich), neomycin (1 g/L, SigmaAldrich), ampicillin (1 g/L, SigmaAldrich), and metronidazole (1 g/L, Alfa Aesar)) sweetened with aspartam (0.05 g/L, Assugrin). Mice were then gavaged six times over the next two weeks with 2 x 10^8^ CFU *E. gallinarum* in PBS with aspartam or PBS with aspartam alone (Control). Three days after the last gavage, mice started receiving a topical treatment of 1.25 mg 5% Imiquimod (IMQ) cream (Aldara 5% Creme, Meda) on the skin of the ear (alternating between treatments) three times a week for 6 weeks, which induces a lupus-like disease in SPF and gnotobiotic settings as previously described (*54*) (*55*).

During the experiment urine and feces were collected weekly, and blood was drawn every two weeks. Splitting cohorts and alternating the blood drawing schedule allowed readouts for every week of treatment. Proteinuria was measured by Albustix reagent strips for urine analysis (Siemens). Serum from blood was extracted using Serum Gel-Z (Sarstedt). *E gallinarum* presence in feces was determined via PCR as described (*18*) (*19*).

Experiments were approved by the Kantonales Veterinäramt BS (Protocol 3042) as well as the Roche animal welfare officer.

### Longitudinal evaluation of *E. gallinarum* translocation in NZW x BXSB F_1_ mice

Male NZW x BXSB F_1_ mice were randomly mixed with littermates from different parental cages at three weeks of age and maintained in an SPF environment as previously described (*18*). Water and a standard laboratory diet were provided ad libitum. Mice were euthanized at 14, 16 and 18 weeks of age, and a laparotomy was performed. Under sterile conditions, mesenteric veins, mesenteric lymph nodes, liver, and spleen were harvested and transferred aseptically to individual BBL Mycoflask Thioglycolate (Fluid) Prepared Media (BD Diagnostic Systems) and incubated for 72 h at 37°C. Next, 100 µL of volume from the BBL Mycoflask Thioglycolate tissue culture was streaked in Gifu Anaerobic Media (GAM) agar plates (HIMEDIA Laboratories) and incubated for 24 h at 37°C. Single colonies were picked and cultivated in GAM broth for 48 h at 37°C, and each culture was collected and pelleted for 10 minutes at 10,000 x *g*. The bacterial pellet was processed using DNeasy Blood and Tissue kit (QIAGEN), and DNA was extracted; DNA was submitted to PCR amplification of the 16S region followed by Sanger sequencing. The sequencing results were analyzed with the SnapGene software, and bacterial identification was assessed with the NCBI BLAST nucleotide database.

Animal care and handling was approved by the Yale Institutional Animal Care and Use Facility and in accordance with the National Institutes of Health (NIH) Guide for the Care and Use of Laboratory Animals.

PCR primers for species-specific detection of *E. gallinarum* in the feces:

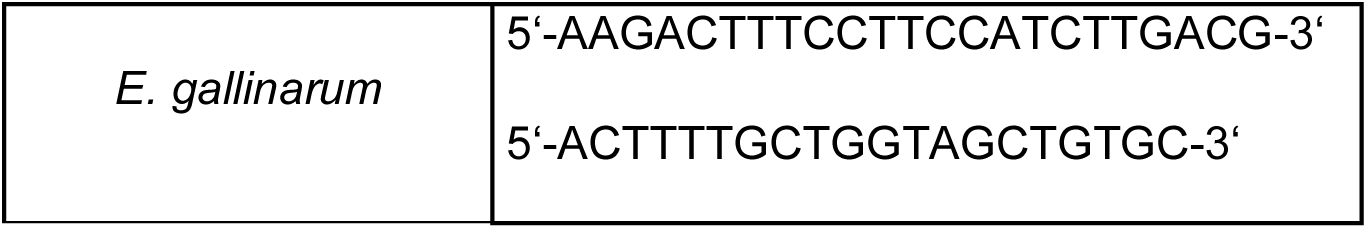

### Statistical Analyses

Statistical parameters including the exact value of n, confidence intervals (mean `±` SD) and statistical significance are reported in the Figures and Figure Legends. Data was considered to be statistically significant when p <0.05 by a Student’s t-test or linear regression model analysis.

* p < 0.05; ** p <0.01; *** p < 0.001. Exact p-values were in addition stated for observed non-significant trends. Statistical analysis was performed in GraphPad Prism.

## Supplementary figure legends

**Fig S1.**
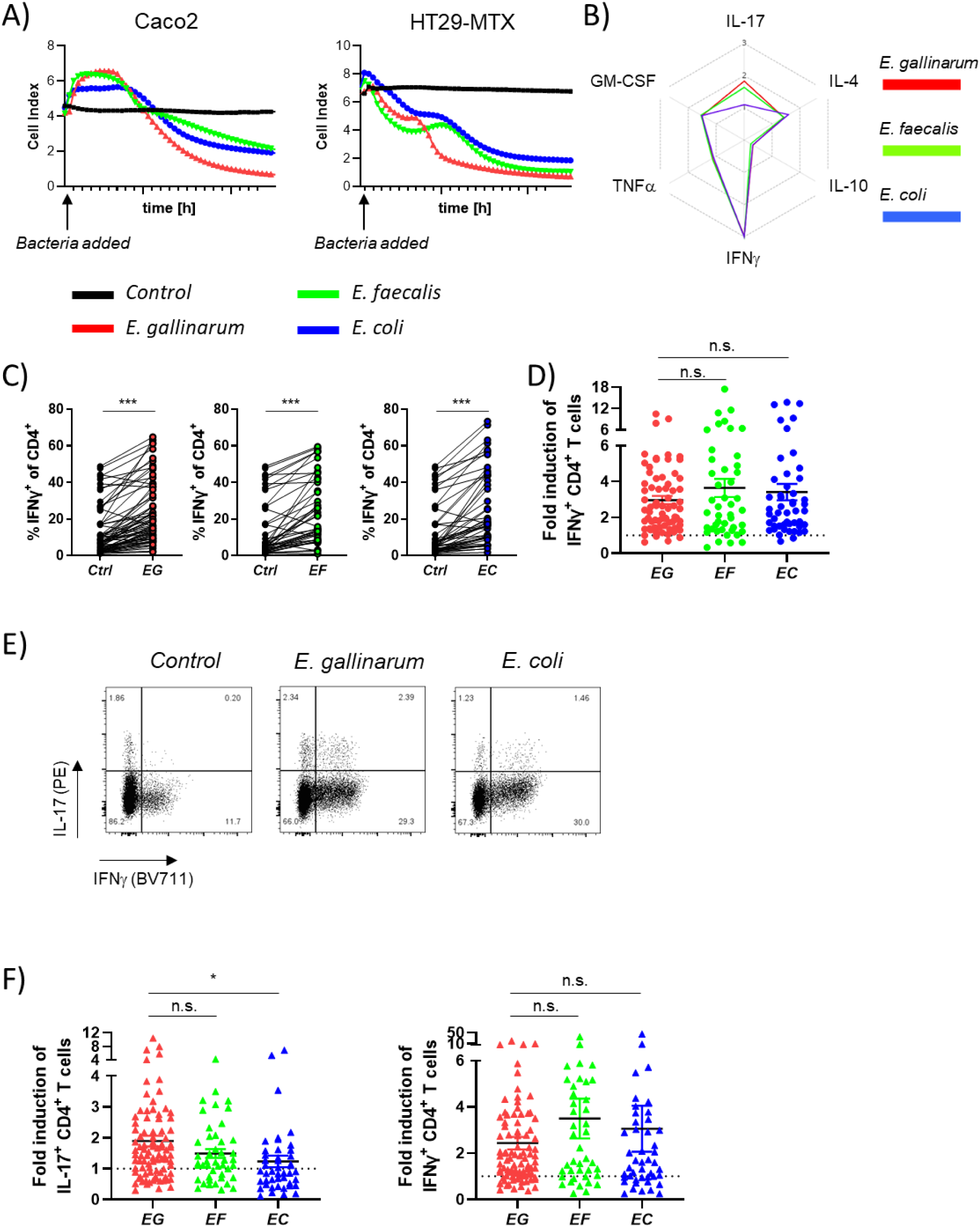
Gut epithelial barrier effects and human Th17/Th1 induction *in vitro* by gut bacteria. a) Epithelial colonic cell lines Caco-2 and HT29-MTX (mucus-producing) were grown to confluency. 10^10^ CFU of *E. gallinarum*, *E. faecalis*, or *E. coli* were added to each well and barrier integrity was monitored in real-time by an xCELLigence cell analyzer. b) Human PBMCs were stimulated with bacterial lysates in the presence of CD3/CD28 activation. Cytokine production by CD4^+^ T cells was assessed by flow cytometry on day 7. Comparison of average fold induction of several T cell-derived cytokines induced by *E. gallinarum (red)*, *E. faecalis (green)*, or *E. coli (blue)* lysates. c) Percentage of IFNγ-expressing CD4^+^ T cells in response to bacterial stimulation. d) Fold induction of IFNγ^+^ CD4^+^ T cells over control by *E. gallinarum (EG)*, *E. faecalis (EF)* or *E. coli (EC)* in PBMCs on day 7 (EG n=69, EF n=45, EC n=49). e) PBMCs were stimulated with bacterial lysates without additional CD3/CD28 activation. Representative plots for no bacterial lysates (*Control*), *E. gallinarum* lysate and *E. coli* lysate. f) Fold induction of IL-17- and IFNγ-producing CD4^+^ T cells by *E. gallinarum (EG)*, *E. faecalis (EF)*, or *E. coli (EC)* in PBMCs on day 7 (EG n=88, EF n=45, EC n=45) without additional CD3/CD28 activation. Statistical analysis: Unpaired t-test (d,f); paired t-test (c)

**Fig S2.**
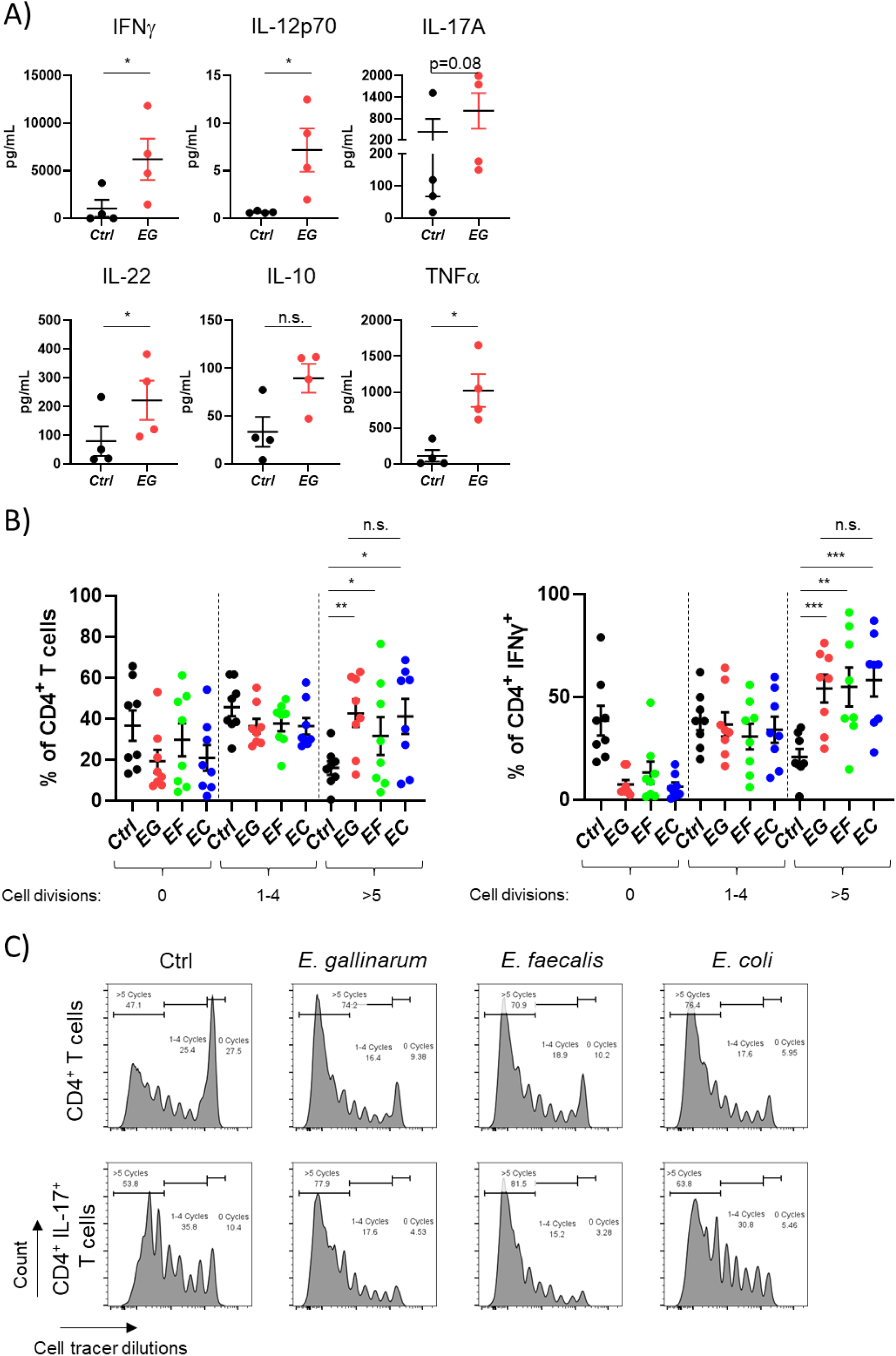
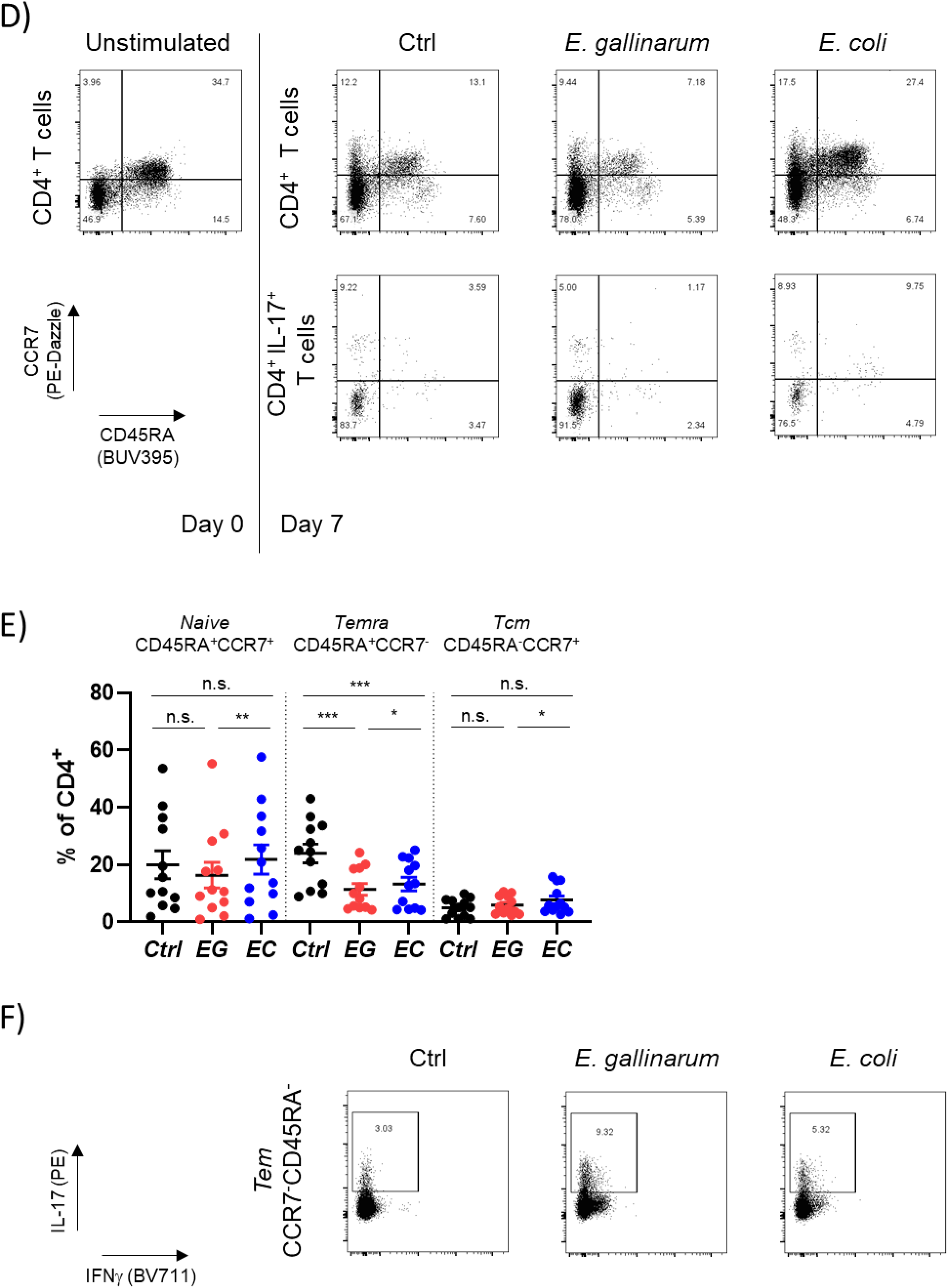
Th17/Th1-related cytokine secretions and frequencies as well as proliferation of human immune cells after stimulation with gut bacterial lysates. Human PBMCs were stimulated with bacterial lysates in the presence of CD3/CD28 activation for 7 days. Cytokines were measured in the supernatant on day 7. a) Secreted proteins were measured in the supernatants by a bead-based multiplex assay after stimulation with *E. gallinarum (EG)* lysate. b) PBMCs were incubated with a cell proliferation dye prior to stimulation with *E. gallinarum (EG)*, *E. faecalis (EF)* or *E. coli (EC)* lysate. The number of cell divisions was assessed after one week. Shown here are divisions for all CD4^+^ T cells (left) and IFNγ^+^ CD4^+^ T cells (right). c) Representative flow cytometry plots for cell tracer analysis for all CD4^+^ T cells and IL-17^+^ CD4^+^ T cells. d) Representative flow cytometry plots for CCR7 and CD45RA for all CD4^+^ T cells and IL-17^+^ CD4^+^ T cells. e) Percentages of CD45RA^+^CCR7^+^ naive, CD45RA^+^CCR7^-^ terminally differentiated (Temra), and CD45RA^-^CCR7^+^ central memory (Tcm) cells of CD4^+^ T cells after stimulation with *E. gallinarum (EG)* or *E. coli (EC)* as measured by flow cytometry. f) Representative flow cytometry plots for IL-17 production by effector memory CD4^+^ T cells after bacterial stimulation. Statistical analysis: paired t-test (b,d), Ratio paired t-test (a)

**Fig S3.**
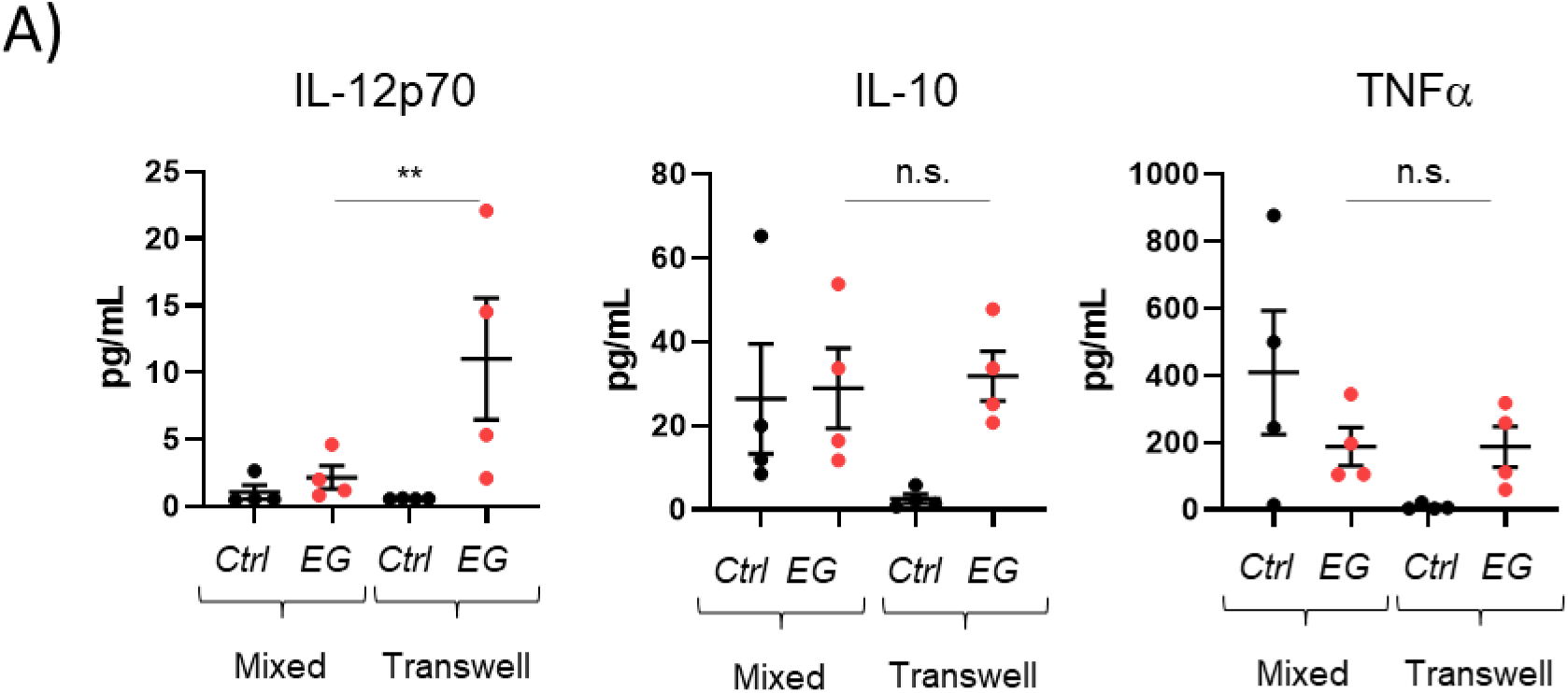
Cytokine production induced by *E. gallinarum* in human CD4^+^ T cell/monocyte co-cultures with or without cell-cell contact *in vitro*. a) Human PBMCs were isolated and CD14^+^ monocytes and CD4^+^ T cells were isolated via magnetic bead sorting. Cells were stimulated with *E. gallinarum (EG)* lysate for 7 days with additional CD3/CD28 activation. CD4^+^ T cells and CD14^+^ monocytes were either mixed or kept separated by a transwell system and stimulated with *E. gallinarum* lysates. Secreted cytokines (IL-12p70, IL-10, TNFα) were measured in the supernatants of cultures by a bead-based multiplex assay. Statistical analysis: Ratio paired t-test (a)

**Fig S4.**
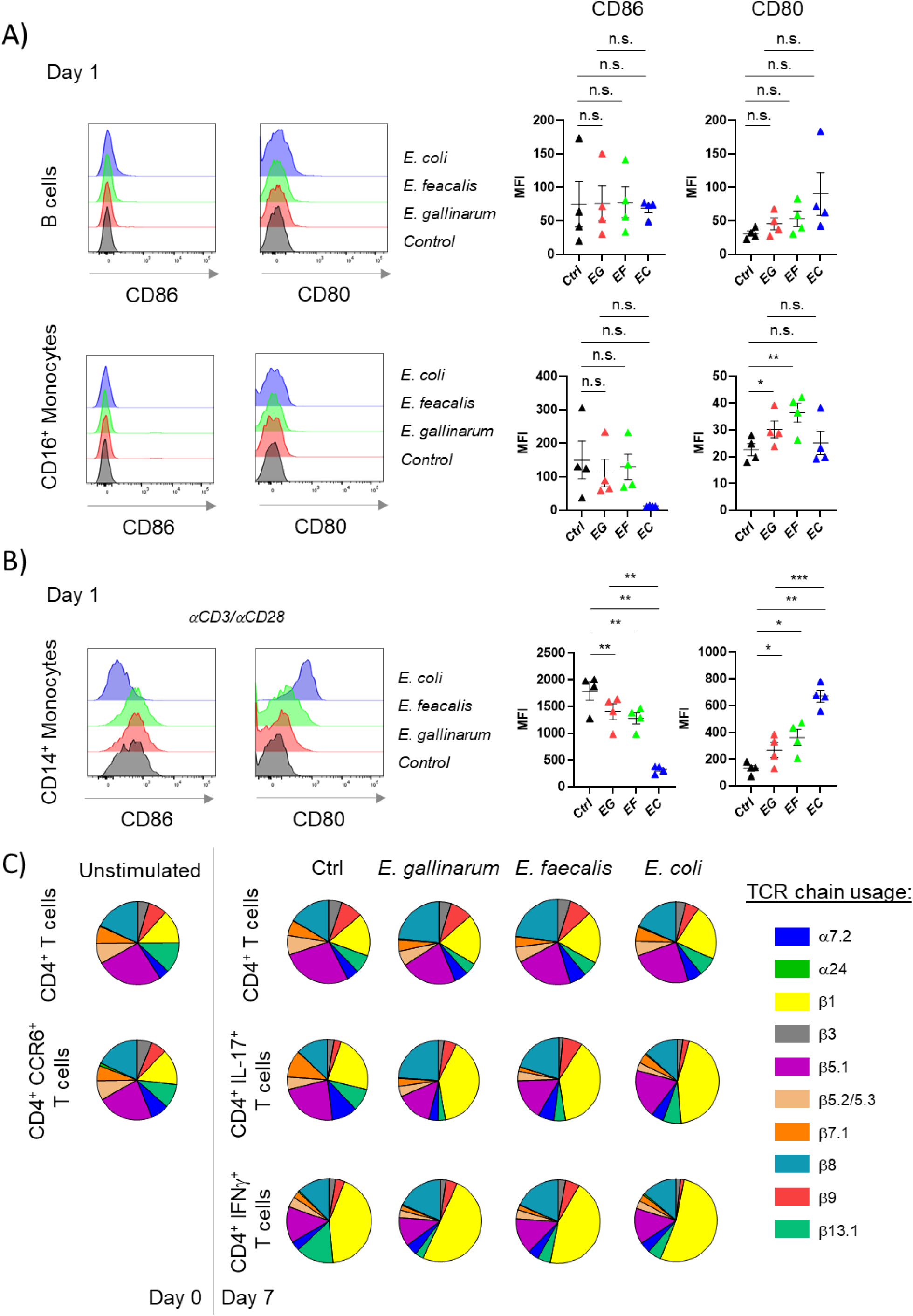
Gut bacterial effects on co-stimulatory molecules on human APCs and on TCR chain distributions in PBMCs. Human PBMCs were stimulated with *E. gallinarum (EG)*, *E. faecalis (EF)*, or *E. coli (EC)* lysates. a) Expression of co-stimulatory molecules CD80 and CD86 by different APCs assessed by flow cytometry on day 1 after stimulation. b) Expression of co-stimulatory molecules CD80 and CD86 by CD14^+^ monocytes in the presence of CD3/CD28 activation assessed by flow cytometry on day 1 after stimulation. c) Expression of different α and β chains of the TCR were assessed by flow cytometry. Circles show distribution of the different chains of one representative donor (of a total of four donors) among CD4^+^ T cells and cytokine-producing subsets on day 0 and on day 7 after stimulation with *E. gallinarum*, *E. faecalis* or *E. coli*. Statistical analysis: paired t-test (a,b)

**Fig S5.**
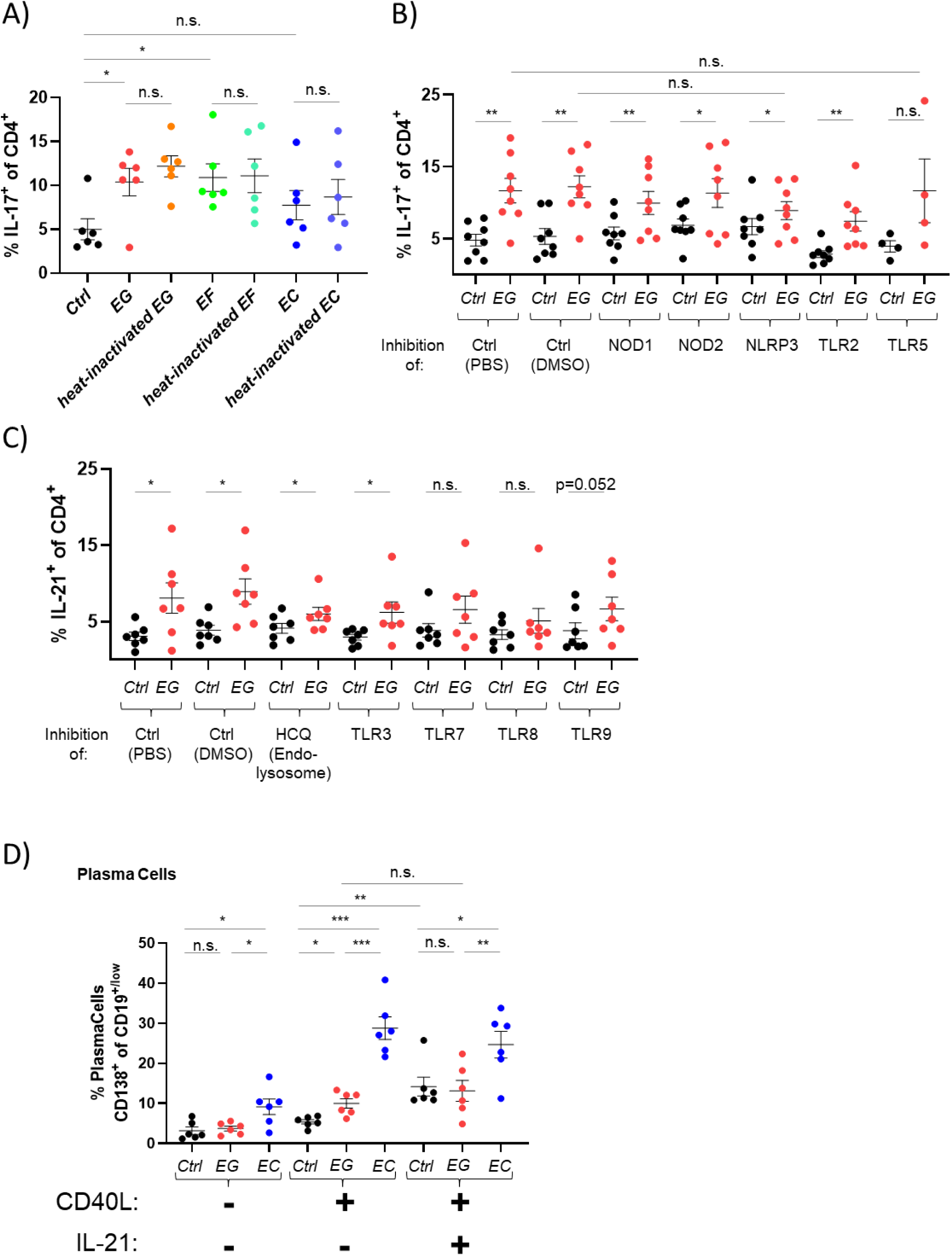
IL-17^+^/IL-21^+^ CD4+ T cell and plasma cell frequencies after *E. gallinarum* stimulation of PBMC with PRR inhibitors or addition of B cell co-stimulatory factors. Human PBMCs were stimulated with bacterial lysates in the presence of CD3/CD28 activation. Cytokine production by CD4^+^ T cells was assessed by flow cytometry on day 7. a) *E. gallinarum (EG)*, *E. faecalis (EF)* or *E. coli (EC)* lysates were either only sonicated or heat-inactivated prior to being used to stimulate PBMCs. Percentage of IL-17^+^ CD4^+^ T cells was measured on day 7 by flow cytometry. b,c) PBMCs were treated with soluble inhibitors of several pattern recognition receptors (b) selective NOD1, NOD2, NLRP3, inhibitors, blocking antibodies against TLR2/TLR5 or c) hydroxychloroquine, TLR3/7/8/9 inhibitors). Cells were subsequently stimulated with *E. gallinarum (EG)* lysate in the presence of CD3/CD28 activation. b) IL-17 and c) IL-21 induction in CD4^+^ T cells was measured on day 7 by flow cytometry. d) In addition to stimulation with *E. gallinarum (EG)* or *E. coli (EC)* lysates, PBMCs were treated with recombinant CD40L and IL-21. Frequencies of plasma cell populations were assessed by flow cytometry on day 7. Statistical analysis: paired t-test (a,b,c,d)

**Fig S6.**
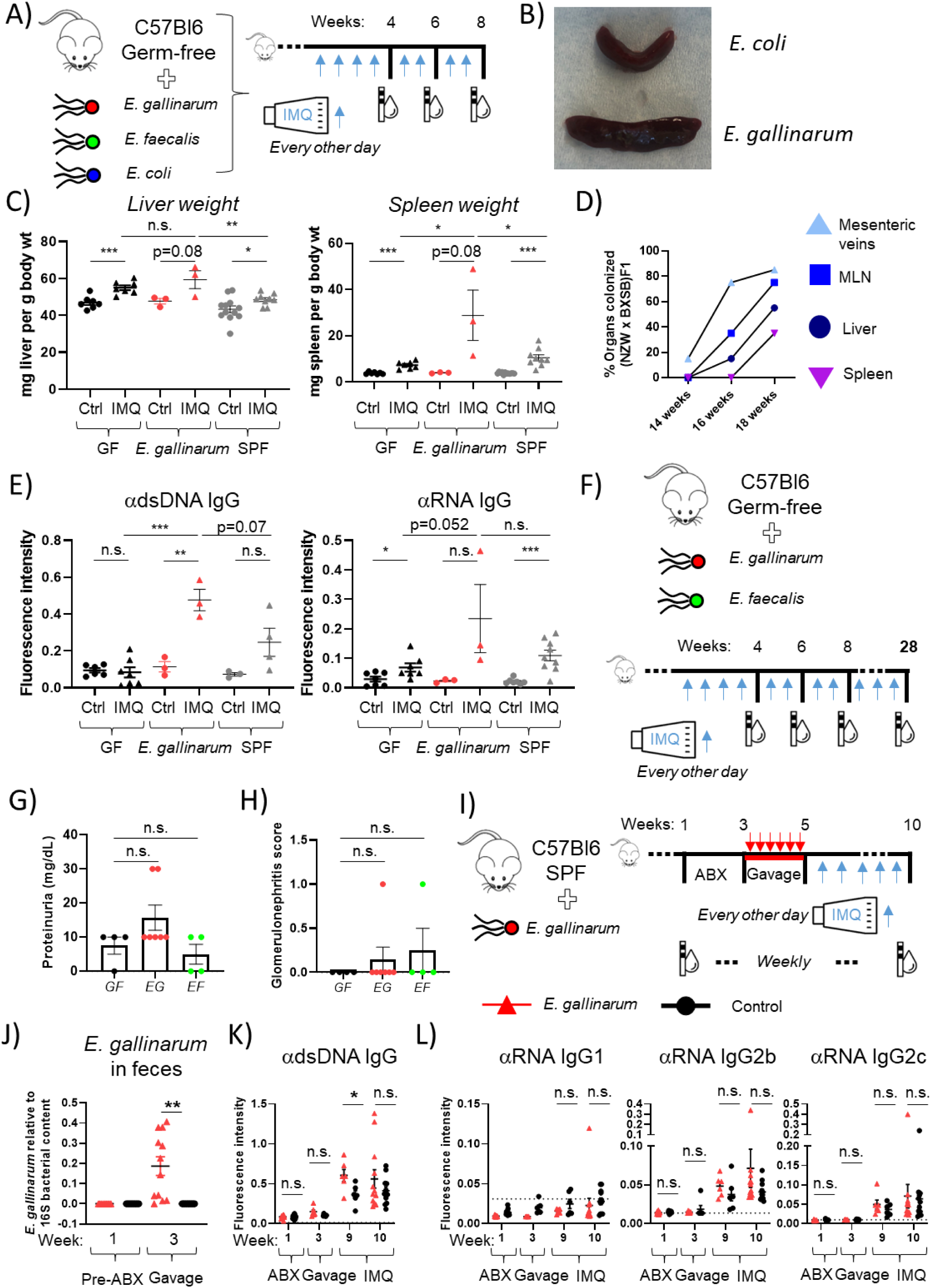
Gut bacterial effects on extraintestinal organ dysfunctions and systemic autoantibody subclasses in gnotobiotic mice, and translocation dynamics of *E. gallinarum* in NZW x BXSB F_1_ mice. a-h) C57BL/6-Germ-free mice were monocolonized with *E. galllinarum (EG)*, *E. faecalis (EF)* or *E. coli (EC)* and treated with topical imiquimod (IMQ). a) Schematic representation of the short-term (8-week) monocolonization studies plus IMQ treatment and blood/urine collection schedules. b) Representative images of the spleens of *E. coli-* or *E. gallinarum*-monocolonized mice after 8 weeks of treatment with IMQ. c) Comparison of relative liver and spleen weight of C57BL/6 germ-free, *E. gallinarum*-monocolonized, and SPF mice d) Temporo-spatial translocation dynamics in NZW x BXSB F_1_ mice naturally colonized by *E. gallinarum*. Mesenteric veins, mesenteric lymph nodes (MLN), livers, and spleens of NZW x BXSB F_1_ mice were harvested at 14, 16, and 18 weeks of age (n = 20 mice each across 4 independent experiments). Tissues were sterilely assessed for bacterial translocation of *E. gallinarum* by culture followed by species-specific PCR. e) Germ-free, *E. gallinarum*-monocolonized, and SPF mice were treated with topical IMQ for 8 weeks. Antibodies targeting dsDNA (left) and RNA (right) were assessed in sera of mice as indicated. f) Schematic representation of long-term (28-week) monocolonization studies plus IMQ treatment and blood/urine collection schedules. g) Proteinuria determined in urine and h) glomerulonephritis score at week 28 of IMQ treatment in germ-free (GF), *E. gallinarum (EG)-* and *E. faecalis (EF)*-monocolonized mice (long-term). i) Schematic representation of introduction of *E. gallinarum* into SPF mice plus IMQ treatment and blood/urine collection schedules. i-l) C57BL/6 SPF mice were treated with oral antibiotics (ABX) and subsequently gavaged with 2×10^8^ CFU of *E. gallinarum* or PBS (Control). After gavage, mice were treated with topical imiquimod every other day. Urine and blood were collected as indicated. Treatment groups were divided into two cohorts, blood was taken every two weeks, alternating between the cohorts and shifting by a week. j) Detection of *E. gallinarum* in the feces of C56BL/6 mice by qPCR before ABX treatment and after *E. gallinarum* gavage. k,l) Serum reactivity for IgG against dsDNA (k) and IgG1, IgG2b and IgG2c against RNA (l) in control- and *E. gallinarum*-gavaged mice before ABX treatment (Week 1), after first gavage (Week 3) or after 6 weeks of IMQ treatment (Week 9 and 10). The dotted line represents the detection threshold. Statistical analysis: Unpaired t-test (c,e,g,h,j,k,l)

**Fig S7.**
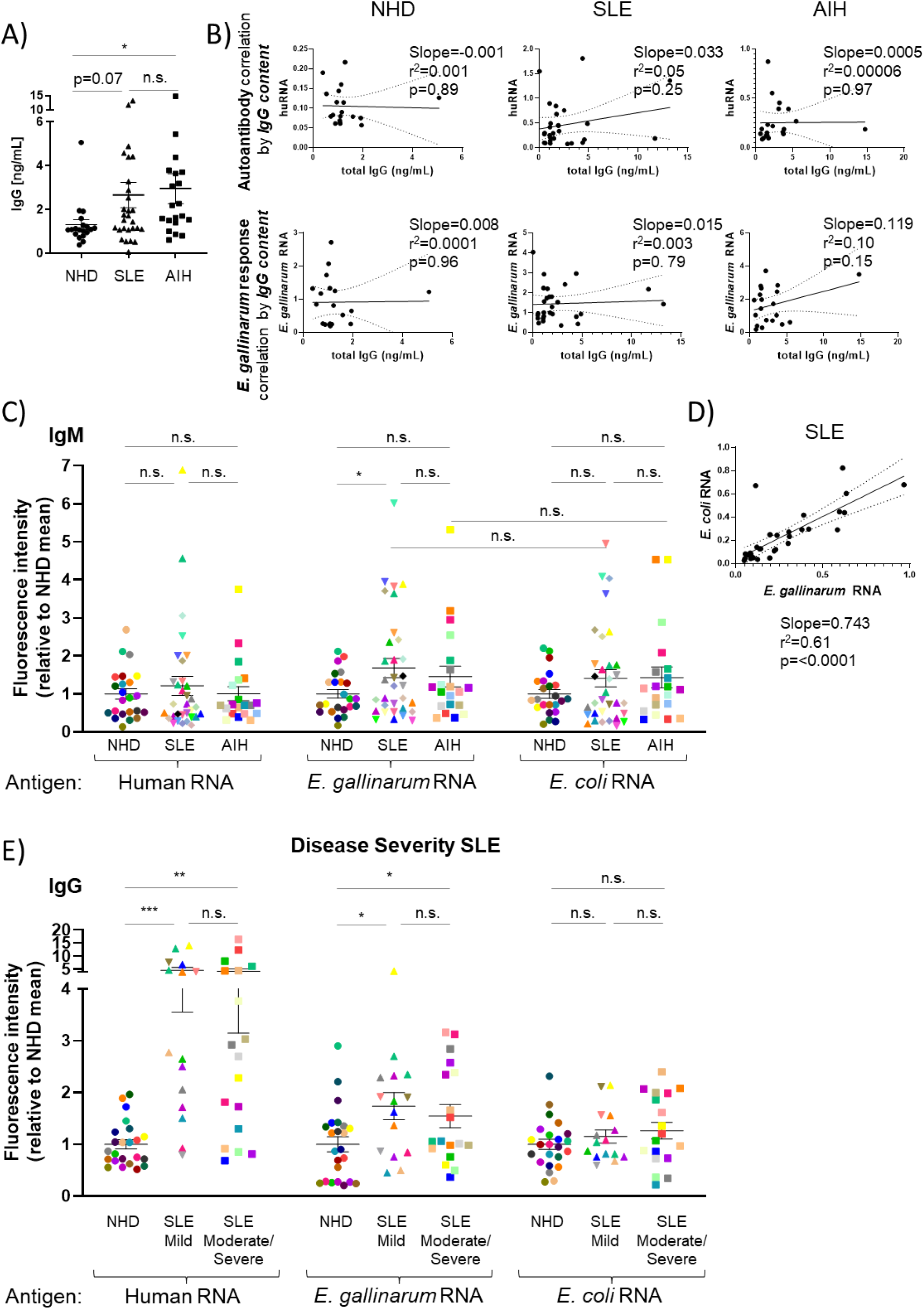
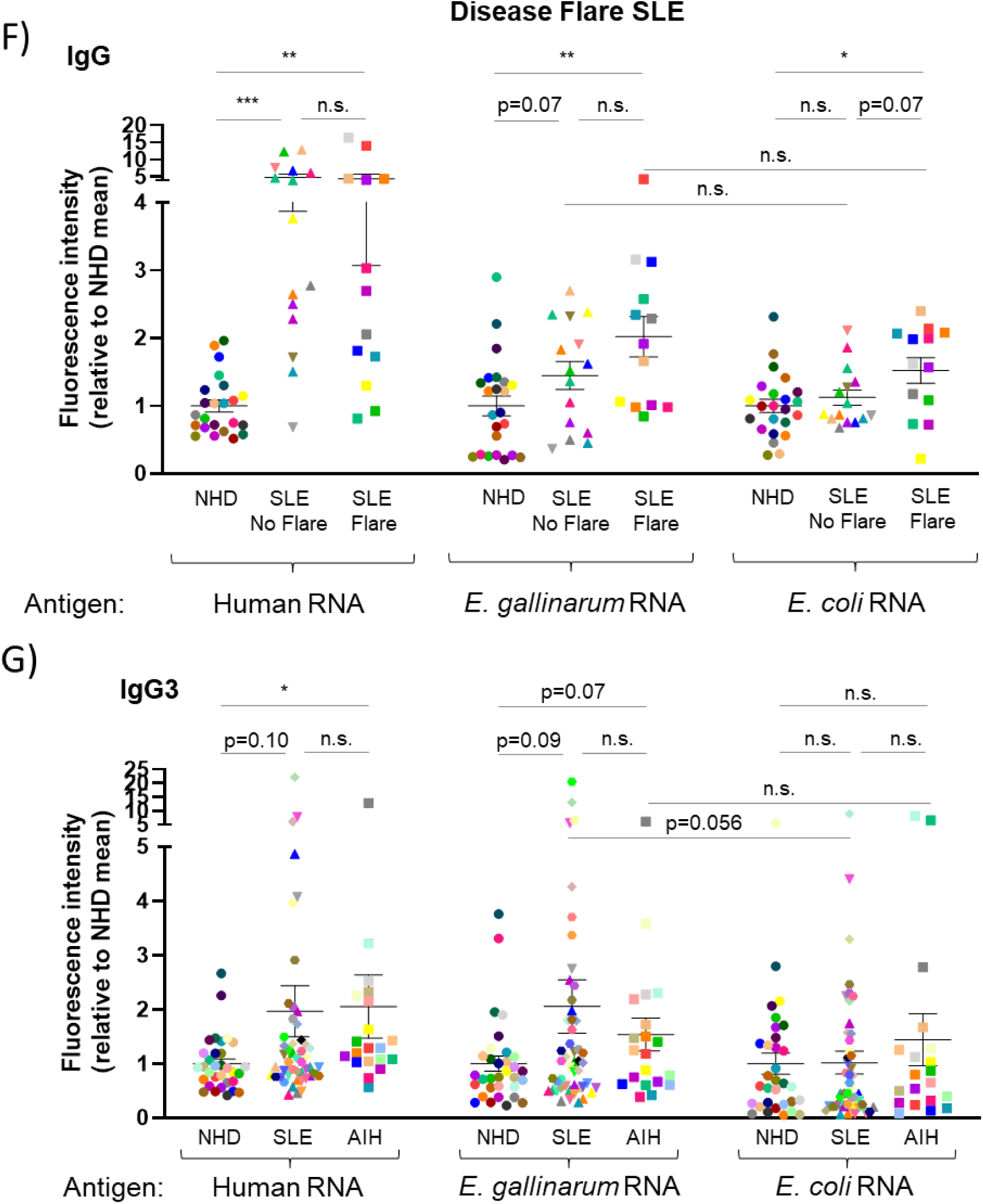
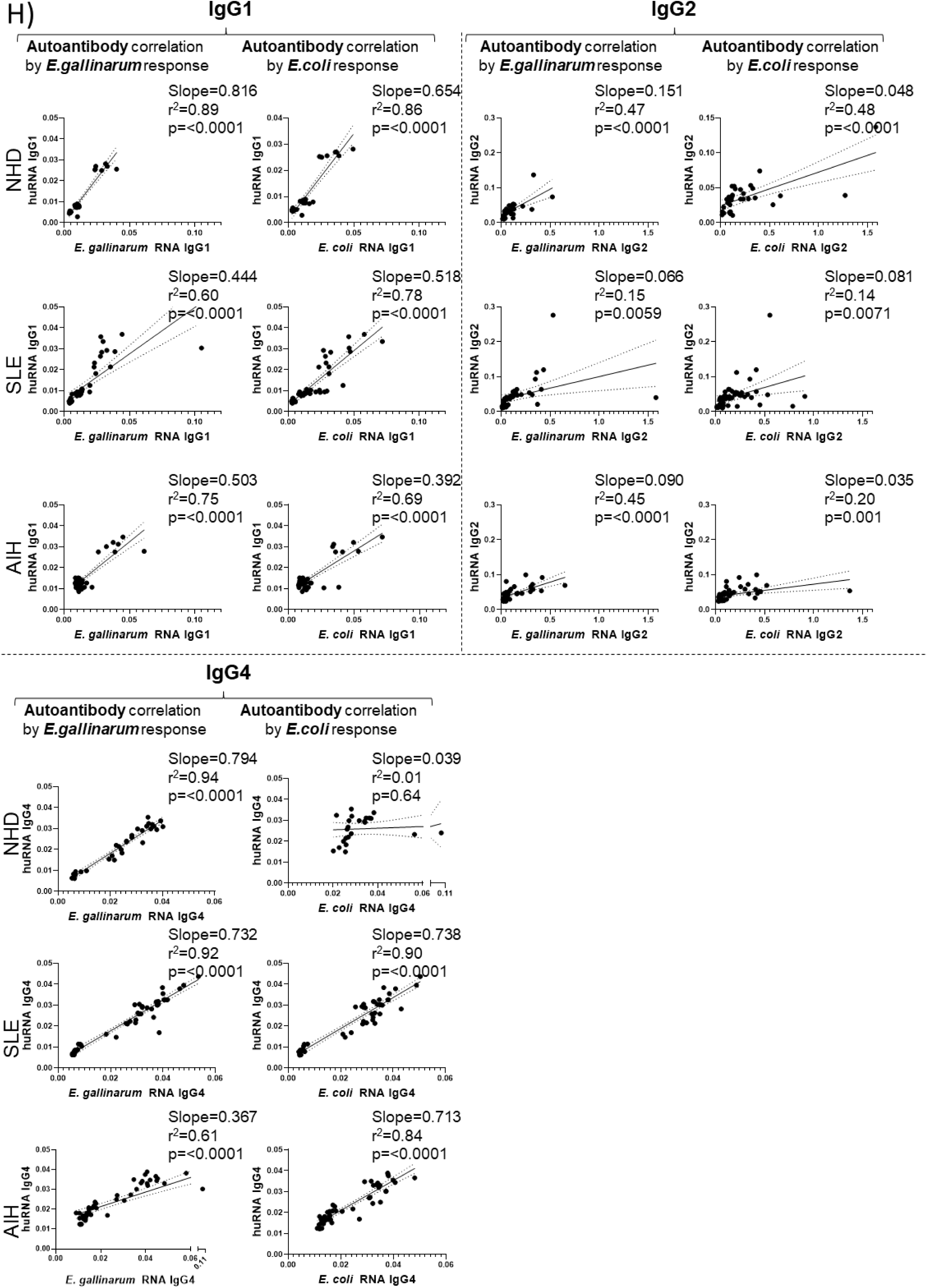
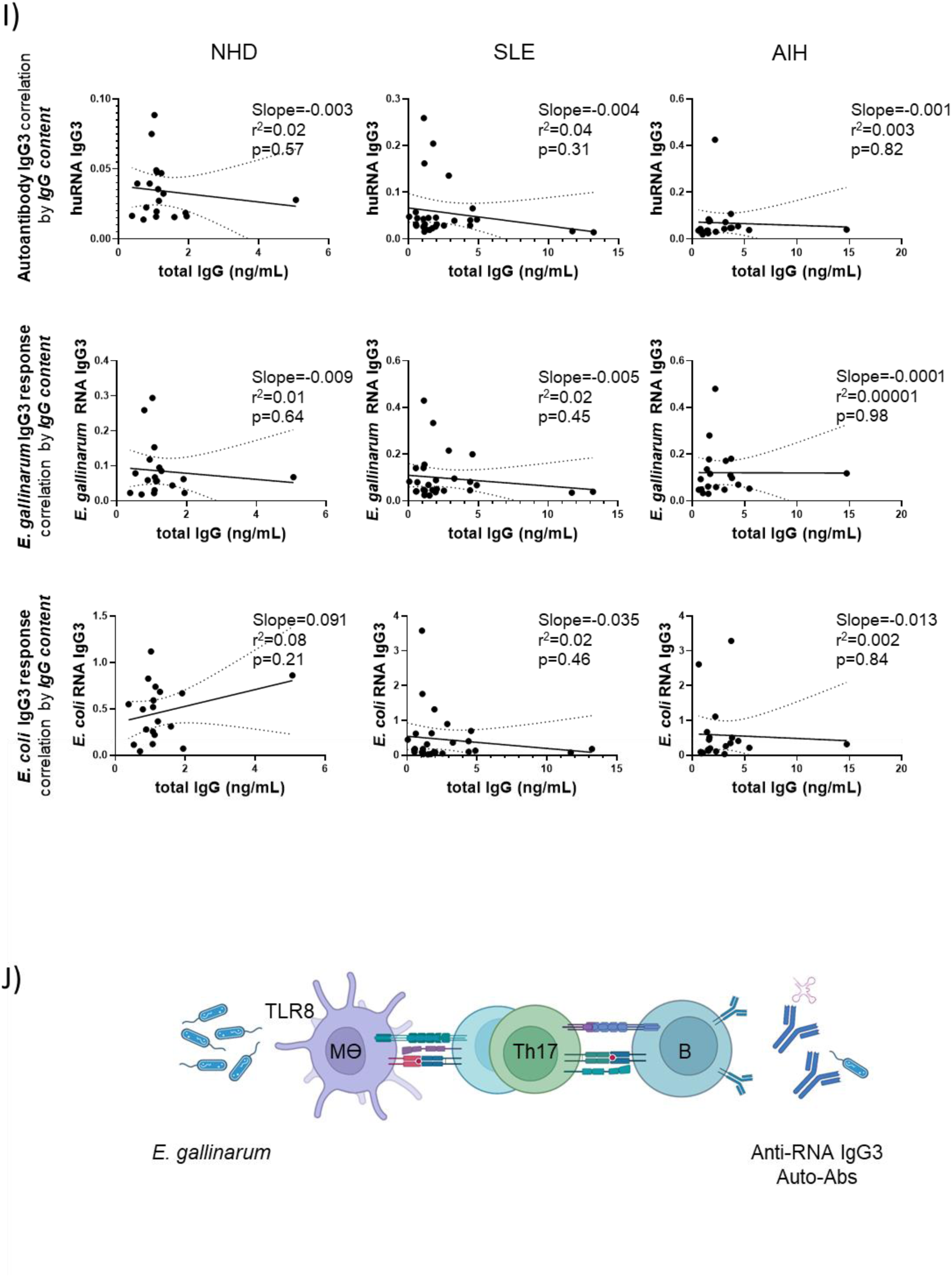
SLE and AIH patient serum reactivities against *E. gallinarum* and *E. coli* RNA. a) Total IgG concentrations in sera of normal healthy donors (NHD), systemic lupus erythematosus patients (SLE), and autoimmune hepatitis patients (AIH), respectively. b) Correlation between total IgG concentration and reactivity against human RNA (upper panel) or *E. gallinarum* RNA (lower panel) for each group (NHD, SLE, AIH). c) Total IgM reactivity was tested in the sera from NHD subjects, SLE, and AIH patients against different RNA preparations. Results were normalized to the average of NHD controls for each RNA preparation. d) Correlation between absolute reactivity of total IgM against *E. coli* RNA and *E. gallinarum* RNA for SLE patients. e) Total IgG reactivity against different RNA preparation was tested in sera from NHD subjects and SLE patients divided based upon their disease severity (mild vs moderate/severe). Results were normalized to the average of NHD controls for each RNA preparation. f) Total IgG reactivity against different RNA preparation was tested in sera from NHD subjects and SLE patients divided based upon their disease flare status (no flare, active/ongoing flare). Results were normalized to the average of NHD controls for each RNA preparation. g) IgG subclass reactivity against different RNA preparation was tested in sera from NHD subjects, SLE and AIH patients, respectively. Results were normalized to the average of NHD controls for each RNA preparation. h) Correlation between absolute reactivity of IgG1, IgG2, and IgG4 against human RNA and *E. gallinarum* RNA (left panels) or *E. coli* RNA (right panels) for each group (NHD, SLE, AIH). i) Correlation between total IgG concentration and IgG3 reactivity against human RNA (upper panel), *E. gallinarum* RNA (middle panel), or *E. coli* RNA (lower panel) for each group (NHD, SLE, AIH). j) Illustration of the proposed model of how translocating *E. gallinarum* may induce extraintestinal Th17 and IgG3 autoantibody responses via its effects on antigen-presenting cells (APCs). *E. gallinarum*-stimulated APCs (monocytes/macrophages) support the differentiation of Th17 cells via T cell-monocyte contact-dependent mechanisms involving monocyte TLR8 signals. Subsequently, Th17 cells may support B cell maturation, which preferentially leads to IgG3 secretion targeting bacterial as well as human RNA-related autoantigens. Image created with Biorender.com. Statistical analysis: Unpaired t-test (a,c,e,f,g); linear regression (b,d,h,i)

